# On-target, dual aminopeptidase inhibition provides cross-species antimalarial activity

**DOI:** 10.1101/2023.10.01.560396

**Authors:** Rebecca C.S. Edgar, Tess R. Malcolm, Ghizal Siddiqui, Carlo Giannangelo, Natalie A. Counihan, Matthew Challis, Sandra Duffy, Mrittika Chowdhury, Jutta Marfurt, Madeline Dans, Grennady Wirjanata, Rintis Noviyanti, Kajal Daware, Chathura D. Suraweera, Ric N Price, Sergio Wittlin, Vicky M. Avery, Nyssa Drinkwater, Susan A. Charman, Darren J. Creek, Tania F. de Koning-Ward, Peter J. Scammells, Sheena McGowan

**Affiliations:** School of Medicine, Deakin University, Geelong, Australia; The Institute for Mental and Physical Health and Clinical Translation, Deakin University, Geelong, Australia; Monash Biomedicine Discovery Institute and Department of Microbiology, Monash University, Clayton, Australia; Drug Delivery, Disposition and Dynamics, Monash Institute of Pharmaceutical Sciences, Monash University, Parkville, Australia; Discovery Biology, Centre for Cellular Phenomics, Griffith University, Nathan, Qld, Australia; Global Health and Tropical Medicine Division, Menzies School of Health Research, Charles Darwin University, Darwin, Northern Territory, Australia; Eijkman Institute for Molecular Biology, Jakarta, Indonesia; Centre for Tropical Medicine and Global Health, Nuffield Department of Clinical Medicine, University of Oxford, Oxford, UK; Mahidol-Oxford Tropical Medicine Research Unit, Faculty of Tropical Medicine, Mahidol University, Bangkok, Thailand; Swiss Tropical and Public Health Institute, Allschwil, Switzerland; University of Basel, Basel, Switzerland; School of Environment and Science, Griffith Sciences, Griffith University, Nathan, Queensland, Australia; Centre for Drug Candidate Optimisation, Monash Institute of Pharmaceutical Sciences, Monash University, Parkville, Australia; Medicinal Chemistry, Monash Institute of Pharmaceutical Sciences, Monash University, Parkville, Australia

**Author notes:** Co-corresponding authors: Tania de Koning Ward; Peter Scammells; Sheena McGowan.

**Keywords:** Malaria, drug resistance, aminopeptidase, Plasmodium falciparum, Plasmodium vivax

## Abstract

To combat the global burden of malaria, development of new drugs to replace or complement current therapies are urgently required. As drug resistance to existing treatments and clinical failures continue to rise, compounds targeting multiple life cycle stages and species need to be developed as a high priority. Here we show that the compound **MMV1557817** is a nanomolar inhibitor of both *Plasmodium falciparum* and *Plasmodium vivax* aminopeptidases M1 and M17, leading to inhibition of end stage haemoglobin digestion in asexual parasites. Multi-stage analysis confirmed that **MMV1557817** can also kill sexual stage *P. falciparum*, while cross-resistance studies confirmed the compound targets a mechanism of action distinct to current drug resistance mechanisms. Analysis of cross reactivity to homologous human enzymes shows the compound exhibits a high level of selectivity, whilst safety as well as druggability was confirmed in the murine model *P. berghei*. **MMV1557817-**resistant *P. falciparum* parasites displayed only low-level resistance (<3-fold) and exhibited a slow growth rate that was quickly outcompeted by wild type parasites. **MMV1557817-**resistant parasites digest significantly more haemoglobin and possess a mutation in *Pf*A-M17 that induces partial destabilisation of the *Pf*A-M17 homohexamer, resulting in high-level resistance to specific *Pf*A-M17 inhibition, but enhanced sensitivity to specific *Pf*A-M1 inhibition, and importantly, these parasites were highly sensitive to artemisinin. Overall, these results confirm **MMV1557817** as a potential lead compound for further drug development and highlight the potential of dual inhibition of M1 and M17 as an effective multi-species drug targeting strategy.

## INTRODUCTION

Malaria remains a leading cause of global mortality, with over 600,000 deaths reported in 2022 arising from infection with *Plasmodium spp*, the majority of these caused by *P. falciparum* and/or *P. vivax* (1). For the past decade, artemisinin-based combination therapies (ACTs) have been recommended by the World Health Organization (WHO) as the standard malaria treatment worldwide. Their widespread use has led to the development of artemisinin resistance in the form of delayed parasite clearance, alongside the rise of partner drug resistance, most notably within South-East Asia and more recently within highly endemic regions of Africa (2–4). Continued emerging resistance jeopardizes the elimination milestones set by the WHO (1), and highlights the urgent need to develop and deploy new antimalarial agents with novel targets and mechanisms of action.

The *Plasmodium* metallo-aminopeptidases M1 and M17 are potential targets worthy of antimalarial development (5–11). These exo-peptidases function at the terminal stages of intra-erythrocytic haemoglobin digestion, a process that occurs in a specialised digestive vacuole (DV) within the parasite and which is required for survival (12). These enzymes utilize a metal-dependent mechanism to facilitate hydrolysis of the N-terminal residue from small peptide substrates (13, 14). The M1 aminopeptidase enzymes have a broad substrate specificity and can cleave N-terminal basic or hydrophobic residues from a peptide substrate (15, 16). In contrast, the *Plasmodium* M17 aminopeptidases display a restricted specificity and generally only act on peptides with an *N*-terminal leucine or tryptophan residue. Although M1 and M17 enzymes have distinct structural and functional features, their overall mechanism of action is the same, with both possessing their own S1 and S1’ substrate binding pockets and essential divalent zinc ion(s) in their catalytic site (17, 18).

Both M1 and M17 appear to be essential for survival in *P. falciparum* parasites, making them attractive potential drug targets. Selective inhibition of *Pf*A-M1 with a specific activity-based probe resulted in swelling of the DV and parasite death (5, 8) and attempts to knockout the gene have not been successful (19, 20). Whilst it was previously reported that inhibition of *Pf*A-M17 with a specific activity-based probe resulted in ring-stage parasite death, this appears to be due to off-target effects, with genetic knockdown of *Pf*A-M17 resulting in parasite death at the later trophozoite stage (7). Importantly, inhibition of *Pf*A-M1 or knockdown of *Pf*A-M17 expression leads to a build-up of undigested short peptide chains that likely originate from haemoglobin, providing confirmation of their function (7, 8). Multiple studies performed in the murine model *Plasmodium chabaudi chabaudi* have also confirmed that inhibition of these aminopeptidases reduces parasite burden, confirming their druggability in an *in vivo* model (5, 21, 22).

Since inhibition of both M1 and M17 leads to parasite death, we developed potent inhibitors that target both enzymes (6, 9, 11). We have previously reported the synthesis and initial characterisation of **MMV1557817** (*N*-(2-(Hydroxyamino)-2-oxo-1-(3′,4′,5′-trifluoro-[1,1′-biphenyl]-4-yl)ethyl)-3,3-dimethylbutanamide). This compound was found to dually inhibit M1 and M17 aminopeptidases from both *P. falciparum* and *P. vivax* in recombinant assays, and exhibited antiplasmodial activity against *P. falciparum* (11). In this current study, we report the preclinical characterisation of **MMV1557817** and investigate its mechanism of action. **MMV1557817** was effective against *P. falciparum* parasites that are resistant to a wide range of other antimalaria drugs as well as sexual state parasites. **MMV1557817** was also effective against *P. falciparum* and *P. vivax* clinical isolates cultured *ex vivo* and showed promising exposure *in vivo* and could effectively clear a *Plasmodium berghei* infection. We confirmed that **MMV1557817** is on-target for both *Pf*A-M1 and *Pf*A-M17 and found that parasites made resistant to **MMV1557817** had a significantly slower growth rate and an increase in haemoglobin digestion. Overall, these results confirm that **MMV1557817** is a candidate for further development as a novel and promising lead compound, whilst also validating M1 and M17 aminopeptidases as suitable cross-species targets for novel drug development.

## METHODS

### Biochemistry and structural biology

#### Enzyme assays

Inhibition of aminopeptidase activity assays were performed against nine aminopeptidases: the M1 alanyl and M17 leucyl aminopeptidases from *P. falciparum (Pf*A-M1*; Pf*A-M17), *P. vivax (Pv-*M1*; Pv-*M17), and *P. berghei* (*Pb*-M1; *Pb*-M17) and three human M1 homologues: LTA4H (OriGene TP307617), ERAP1 (OriGene TP314469) and ERAP2 (Creative BioMart ERAP2-304H). The *Plasmodium* enzymes were produced recombinantly as described previously (16, 17) and the human recombinant enzymes were purchased from commercial suppliers as indicated.

The ability of **MMV1557817** to inhibit aminopeptidase activity was assessed by fluorescence assays using fluorogenic substrates *L*-Leucine-7-amido-4-methylcoumarin hydrochloride (H-Leu-NHMec) for all enzymes except LTA4H, which was assessed using *L*-Alanine-7-amido-4-methylcoumarin hydrochloride (H-Ala-NHMec, 20 μM). The concentration of Leu-NHMec in the assay was held constant for each enzyme and ranged from 10 – 100 μM, depending on the enzyme. Reactions were measured at 37°C in white 384–well plates in a total volume of 25 or 50 μL using a spectrofluorimeter (BMG FLUOstar) with excitation at 355 nm and emission at 460 nm. The fluorescence signal was continuously monitored until a final steady state velocity, *v*, was obtained. Inhibition constants were calculated from biological triplicates made from three different protein preparations.

For determination of the Morrison inhibition constant (*K*i^app^), enzymes were pre-incubated in 100 mM Tris–HCl, pH 8.0 (supplemented with 1 mM CoCl2 for M17 aminopeptidases) and the inhibitors added 20 min prior to the addition of substrate. Substrate concentrations were selected to allow sensitive detection of enzyme activity while not exceeding the *K*m for each enzyme. A compound concentration range was selected to obtain a complete inhibition curve (0% – 100 %) in biological triplicate. Where possible, the *K*i values were calculated by plotting the initial rates versus inhibitor concentration, and fitting to the Morrison equation for tight-binding inhibitors in GraphPad Prism (non-linear regression method). Where a *K*i^app^ could not be calculated, the percentage inhibition was calculated assuming 100% activity in the absence of compound.

#### Safety pharmacology screening

**MMV1557817** was screened at 10 μM for % inhibition against known human DNA repair (including HDAC1, 2, 5, 7, 8, 9, 10 and sirtuin 3 and 6) and matrix metalloprotease (MMP2, 3, 7, 8, 9, 13 and MT1) enzymes (by Eurofins Cerep, study number 100048667 and 100048668). Results showing an inhibition or stimulation higher than 50% are considered to represent significant effects of the test compounds.

#### Protein crystallography

*Pf*A-M1 and *Pf*A-M17 were co-crystallized with **MMV1557817** or crystalized unbound by the hanging-drop method, using previously established protocols (17, 18). Purified protein was concentrated to 5.0 mg/mL and 15 mg/mL respectively and for co-crystals, mixed with **MMV1557817** to a final ligand concentration of 1 mM. For *Pf*A-M1, crystals grew in 20−30% poly(ethylene glycol) (PEG)8000, 0.1 M Tris pH 7.5−8.5, 0.2 M MgCl2, 10% glycerol. For PfA-M17, crystals grew in 30 – 40 % PEG(400), 0.1 M Tris pH 7.5 – 8.5, 0.2 M Li2SO4. Co-crystals were subjected to an additional overnight compound soak (mother liquor supplemented with 1 mM ligand and 1 mM Zn^2+^ for *Pf*A-M17) before being snap-frozen in liquid nitrogen. Data were collected at 100 K using synchrotron radiation at sthe Australian Synchrotron beamlines 3BM1(23) and 3ID1. Data were processed using iMosflm (24) or XDS (25) and Aimless (26) as part of the CCP4i program suite (27). The structures were solved by molecular replacement in Phaser (28) using the structure of unliganded enzymes (RCSB ID 3EBG for *Pf*A-M1 and 3KQZ for *Pf*A-M17) as the search models. The structures were refined using Phenix (29), with 5% of reflections set aside for calculation of *R*free. Between refinement cycles, the protein structure, solvent, and inhibitors were manually built into 2*F*o − *F*c and *F*o − *F*c electron density maps using COOT (30, 31), with restraint files generated by Phenix where necessary. The data collection and refinement statistics can be found in Supplementary Table 1. The coordinates and structure factors are available from the Protein Data Bank with PDB Accession codes: 8SVL (*Pf*A-M1-**MMV1557817**), 8SVM (*Pf*A-M17-**MMV1557817**); 8SW9 (*Pf*A-M17(A460S)).

### Parasitology

#### Parasite cultures

*P. falciparum* asexual blood stage parasites were cultured in human O^+^ RBCs at 3–5% haematocrit in RPMI-1640 media supplemented with 25 mM HEPES, 50 µM hypoxanthine, 2 mM L-glutamine, 2 g/L sodium bicarbonate, 0.5% (w/v) AlbuMAXII (Invitrogen) and 10 µg/mL gentamycin, in modular incubator chambers at 5% O2, 1% CO2 and 90% N2 at 37°C unless otherwise stated.

#### Determination of MMV1557817 IC50 in laboratory conditions against asexual stage parasites

Viability assays were performed as previously described (7). Sorbitol synchronised *Pf*3D7 at ring stages were cultured in the presence of drug serially diluted across a 96-well plate at 0.5% parasitaemia and 2% haematocrit. Cultures were grown for 72 h before being placed at −80°C. After thawing, cultures were incubated with equal volumes of lysis buffer (20 mM Tris pH 7.5, 5 mM EDTA, 0.008% saponin (w/v) and 0.008% Triton x-100 (v/v)) containing 0.2 μL/mL SYBR Green I Nucleic Acid Gel Stain (10,000× in DMSO; ThermoFisher). After 1 h incubation, fluorescence intensity was read on a Glomax® Explorer Fully Loaded (Promega) at emission wavelengths of 500-550 nm and an excitation wavelength of 475 nm. Graphs were generated using GraphPad Prism (V.8.4.1). Parasite survival was compared to vehicle treated cultures in 3 biological experimental replicates performed in triplicate. This IC50 was used in asexual *P. falciparum* assays unless otherwise stated.

#### Determination of MMV1557817 IC_50_ against sexual stage parasites

Viability assays were performed as described previously (32, 33) against *Pf*NF54-*s16*-GFP early (I-III); late (IV-V) and mature (V) stage gametocytes. Gametocytes were assessed at the appropriate time points following gametogenesis; early stage (day 2), late stage (day 8) and mature (day 12). Compounds diluted in 4% DMSO were transferred into 384-well imaging plates; gametocytes prepared as described previously (35) were added, and plates incubated for 72 h in 5% CO2, 5% O2 and 60% humidity at 37°C. After 72 h incubation, 5 µL of MitoTracker Red CMH2XRos in phosphate buffered saline (PBS) was added per well and plates incubated overnight. Image acquisition and analysis was undertaken on the Opera QEHS micro-plate confocal imaging system. An Acapella based script using the CMH2XRos fluorescent signal and the GFP designated object quantifying viable stage dependent parasite morphology identified gametocytes. Gametocyte viability was calculated as a percentage of the positive (5 µM puromycin) and negative (0.4% DMSO) controls. IC_50_ values were calculated using a 4-parameter log dose, non-linear regression analysis, with sigmoidal dose response (variable slope) curve fit using Graph Pad Prism (ver 4.0). No constraints were used in the curve fit. Chloroquine, artesunate, pyronaridine, pyrimethamine, DHA and Methylene Blue were used as control compounds. Experiments were performed in duplicate for 2 or 3 biological replicates.

#### MMV1557817 IC_50_ against drug resistant field isolates and laboratory selected *P. falciparum* parasites

*In vitro* testing was performed with the modified [^3^H]-hypoxanthine incorporation assay, as previously reported (34).

#### Parasite killing rate assay

Assays were performed as previously described (7). *Pf*3D7 parasite cultures at ring or trophozoite stage were set up at 0.5% parasitaemia and 2% haematocrit in a 96-well plate and treated with 10× IC_50_ of **MMV1557817** or artesunate for 24 or 48 h. Parasite cultures incubated for 48 h were fed at 24 h with fresh media containing compound. The compound was then washed out with 3× washes and cultures were split 1/3 before being allowed to grow for a further 48 h, after which time the plates were placed at −80°C. Once thawed, cultures were analysed using SYBR Green I as described above. Parasite viability was determined as a percentage of vehicle-treated controls, and experiments were performed in 4 biological replicates.

#### Parasite reduction ratio

Assays were performed as described previously with some alterations (35). *Pf*3D7 parasite cultures at 0.5% parasitaemia and 2% haematocrit were treated with 10× the IC_50_ of **MMV1557817** or chloroquine in 96-well plates. Specifically, media containing compound was placed in wells every 24 h for up 120 h. After each of the 5 treatment days (0 hrs, 24 hrs, 48 hrs, 72 hrs, 96 hrs or 120 hrs), a subset of wells were removed and the drug was washed off with 3× washes with culture medium, after which the parasites were aliquoted into a new 96-well plate before being serially diluted 1/3. The cultures were then maintained for a further 3 weeks, during which time the parasites were fed with complete culture medium 3 times a week. Parasite growth was measured at day 21 using SYBR green I as described above. Fluorescence was used to determine at what treatment day wells became positive for growth when compared to the well before it; greater than double the fluorescence was determined to be a positive well. This data was then transformed into a log (parasite viability) +1 value for each day and plotted on GraphPad Prism. Experiments were performed in 2 or 3 biological replicates in quadruplicate.

#### *Ex-vivo* activity of MMV1557817 against drug resistant field isolates

*Plasmodium* isolates were collected from patients attending public health clinics in Timika (Papua, Indonesia), a region endemic for multidrug-resistant strains of *P. vivax* and *P. falciparum* (36–38). Patients with symptomatic malaria presenting to an outpatient facility were recruited into the study if infected with either *P. falciparum* or *P. vivax*, with a parasitaemia of between 2,000 μl^-1^ and 80,000 μl^-1^, and the majority (>60%) of asexual parasites at ring stage of development. Venous blood (5 mL) was collected by venipuncture and after removal of host white blood cells using Plasmodipur filters (EuroProxima B.V., The Netherlands), packed infected red blood cells (iRBCs) were used for the *ex vivo* drug susceptibility assay.

Anti-malarial drugs chloroquine (CQ), piperaquine (PIP), mefloquine (MFQ), and artesunate (AS) (WWARN QA/QC Reference Material Programme), and **MMV1557817** were prepared as 1 mg/mL stock solutions in H2O or DMSO. Drug plates were pre-dosed by diluting the compounds in 50% methanol followed by lyophilization and storage at 4°C. Drug susceptibility of *P. vivax* and *P. falciparum* isolates was measured using a protocol modified from the WHO microtest as described previously (38, 39). Briefly, 200 μL of a 2% haematocrit Blood Media Mixture (BMM), consisting of RPMI 1640 medium plus 10% AB+ human serum (*P. falciparum*) or McCoy’s 5A medium plus 20% AB+ human serum (*P. vivax*) was added to each well of pre-dosed drug plates containing 11 serial concentrations (2-fold dilutions) of the anti-malarials; maximum concentrations shown in brackets: CQ (2,993 nM), PIP (1,029 nM), MFQ (338 nM), AS (49 nM), and **MMV1557817** (356 nM). A candle jar was used to mature the parasites at 37.0°C for 35-56 h. Incubation was stopped when >40% of ring stage parasites had reached the mature schizont stage in the drug-free control wells as determined by light microscopy.

Thick blood films made from each well were stained with 5% Giemsa solution for 30 min and examined microscopically. The number of schizonts per 200 asexual stage parasites was determined for each drug concentration and normalized to the control well without drug. The dose-response data were analysed using nonlinear regression analysis and the IC_50_ value derived using an inhibitory sigmoid Emax model (In Vitro Analysis and Reporting Tool; IVART (40, 41). *Ex vivo* IC_50_ data were only used from predicted curves where the Emax and E0 were within 15% of 100 and 1, respectively.

Ethical approval for this study was obtained from the Eijkman Institute Research Ethics Commission, Eijkman Institute for Molecular Biology, Jakarta, Indonesia (EIREC-47), and the Human Research Ethics Committee of the Northern Territory (NT) Department of Health & Families and Menzies School of Health Research, Darwin, Australia (HREC 2010-1396).

#### Determination of MMV1557817 and artemisinin interaction *in vitro*

Fractional inhibitory concentrations (FIC) were determined as previously described (42). The IC_50_ values were calculated from fixed mixed ratios (5:0 to 0:5) of artemisinin (starting dilution 100 nM) and **MMV1557817** (starting dilution 1000 nM) as described above. The FIC for each drug was determined by IC_50_ of drug in mixture / IC_50_ of drug alone, and sum of FIC was determined by addition of the two individual FIC values. Synergistic FIC is described as < 1, additive = 1 < 2 and antagonism > 2 (42).

#### *In vitro* ADME and *in vivo* exposure in mice

Preliminary studies were conducted to assess the *in vitro* absorption, distribution, metabolism, and excretion (ADME) properties and *in vivo* mouse exposure of **MMV1557817** following single oral dosing. Kinetic solubility was conducted using nephelometry based on a modification of a previously published method (43). Briefly, compound was dissolved in DMSO and spiked into either pH 6.5 phosphate buffer (to reflect the pH of the fasted state upper small intestine) or 0.01 M HCl (to reflect gastric pH) with a final DMSO concentration of 1% (v/v) and final concentrations ranging from 1 to 100 µg/mL. Samples were analysed by nephelometry to determine the concentration above which precipitation occurred.

Metabolic stability was assessed by incubating **MMV1557817** (1 µM) with human, rat or mouse liver microsomes (0.4 mg/mL) over 60 min at 37°C in the absence or presence of an NADPH-regenerating buffer. Samples were collected over the incubation period, quenched by the addition of an equal volume of acetonitrile, and analysed using a Waters Acquity UPLC and Xevo G2 QTOF mass spectrometer with positive electrospray ionization under MS^E^ mode. The natural log of the substrate concentration was plotted against the incubation time to determine the first-order degradation rate constant which was normalized to the microsomal protein concentration to give the *in vitro* intrinsic clearance (*in vitro* CLint, µL/min/mg microsomal protein). Additional metabolism (1 µM substrate) and metabolite identification (10 µM substrate) studies were conducted by incubating **MMV1557817** with human or rat cryopreserved hepatocytes (1.4 x 10^6^ viable cells/mL) suspended in pH 7.4 Krebs-Henseleit buffer over 120 min at 37°C/7.5% CO2. Cell viability was determined by Trypan Blue exclusion. At various times over the incubation period, samples were quenched with the addition of an equal volume of acetonitrile and analysed as described above. Metabolite identification was conducted by accurate mass and MS/MS analysis.

The stability of **MMV1557817** was further assessed by incubating with mouse (purchased from the Animal Resources Centre, Perth, Western Australia) or human (Australian Red Cross Blood Service) plasma over 4 h at 37°C. Plasma was spiked with **MMV1557817** (∼500 ng/mL) and samples were taken periodically, and proteins precipitated with a 2-fold excess of acetonitrile. After centrifugation, the supernatant was collected and stored at −80°C until analysis. Sample analysis was conducted using a Waters Acquity UPLC and Quattro Premier mass spectrometer operated in positive electrospray ionization mode with multiple-reaction monitoring (transition (m/z) 395.27 > 362.28, cone voltage 20 V, CID 7 V). The column was a Supelco Ascentis Express RP Amide (50×2.1 mmm 2.7 µm) and the mobile phase was a water/acetonitrile gradient containing 0.05% formic acid with a 4 min cycle time. The injection volume was 3 µL and the flow rate was 0.4 mL/min. Concentrations were determined by comparison to a calibration curve prepared in blank plasma and processed in the same way as for the samples.

Plasma protein binding (human, rat and mouse plasma) and binding to the Albumax medium used for *in vitro* activity assessment was measured by ultracentrifugation based on a method reported previously (44) under conditions that maintained the pH at 7.4 ± 0.1. Human plasma was obtained from the Volunteer Blood Donor Registry (Walter & Eliza Hall Institute, Melbourne, Australia), rat plasma was from male Sprague Dawley rats (Animal Resource Centre, Perth, Western Australia) and mouse plasma was procured as described above. RPMI-1640 media supplemented with 25 mM HEPES, 2 g/L sodium bicarbonate and 100 mg/L neomycin was supplemented with 5 g/L AlbuMAXII on the day prior to binding assessment. Plasma protein binding was measured using 10% plasma diluted with pH 7.4 phosphate buffered saline, and the extent of binding in neat plasma was then calculated via an established approach which accounts for the shift in equilibria that occurs with protein dilution (45). Each medium was spiked with **MMV1557817** (∼1000 ng/mL for plasma, 1 µM for Albumax), briefly mixed and equilibrated for ∼30 min (Albumax equilibration was conducted in a 5% CO2 incubator to maintain pH 7.4) after which they were subjected to ultracentrifugation at 37°C using a sealed rotor (Beckman Rotor type 42.2 Ti; 223,000 x g) for 4.2 h to separate proteins. Additional aliquots of spiked matrix were maintained at 37°C for 4.2 h but not centrifuged to serve as controls for stability assessment and to obtain a measure of the total concentration in each matrix. Following centrifugation, aliquots of the protein free supernatant and the non-centrifuged matrix controls were ‘matrix matched’ by addition of an equal volume of the opposite medium (i.e. either blank buffer or blank matrix) and assayed as described above with comparison to calibration standards prepared in 50% matrix/50% phosphate buffer.

*In vivo* exposure of **MMV1557817** was assessed in non-infected male Swiss outbred mice in parallel to *in vivo* efficacy studies. All animal studies were conducted using established procedures in accordance with the Australian Code of Practice for the Care and Use of Animals for Scientific Purposes, and the study protocols were reviewed and approved by the Monash Institute of Pharmaceutical Sciences Animal Ethics Committee. **MMV1557817** was dissolved in 70% (v/v) Tween 80/30% (v/v) ethanol and diluted 10-fold with water just prior to dosing. This produced a uniform off-white milky suspension of pH 6.0. **MMV1557817** (50 mg/kg) was dosed by oral gavage (10 mL/kg) to non-fasted mice (29-35 g) and blood samples were collected into heparinized tubes for up to 24 h post-dosing (n=2 mice per time point) with a maximum of two samples per mouse. Samples were collected by either submandibular bleed (∼120 µL) or terminal cardiac puncture (under inhaled isoflurane anaesthesia). Blood samples were centrifuged immediately, and supernatant plasma removed and stored at −80°C until analysis as described above.

#### Efficacy testing of MMV1557817 i*n vivo*

Stock solutions of compounds (**MMV1557817** and artesunate) were dissolved in 70% (v/v) Tween 80/30% (v/v) ethanol and diluted 10-fold with water prior to dosing. Biological assessment of the *in vivo* antimalaria efficacy of **MMV1557817** was assessed using the *P. berghei* rodent malaria 4-day suppressive test (46). Female Balb/c mice at 6-weeks of age in groups of 4-5 were infected intraperitoneally with 2 × 10^7^ *P. berghei* ANKA*-*infected erythrocytes. At 2 h and days 1, 2, and 3 post-infection, mice were orally-gavaged with either 50 mg/kg **MMV1557817**, 30 mg/kg artesunate (Sigma) or vehicle control. Parasitaemia was assessed by visualizing Giemsa-stained thin blood smears by microscopy and a minimum of 1000 RBCs were counted. To calculate percent antimalarial activity the following formula was used: 100 − (mean parasitemia treated/mean parasitemia vehicle control) ×100. These studies were conducted using established procedures in accordance with the Australian Code of Practice for the Care and Use of Animals for Scientific Purposes, and the study protocols were reviewed and approved by the Deakin University Animal Ethics Committee.

#### CRISPR-Cas9 editing of *pfa-m17*

Introduction of the A460S mutation was attempted using methods previously described (47). The CRISPR guide 5’ – AATGGTAAAACTATAGAAGT was ligated into the *Bbs*I site of the plasmid pDC2-cam-Cas9-U6-hDHFR (47). Additionally, the last 1373 base pairs from *pfa-m17* containing the A460S mutation (gct → tct) was synthesised by Integrated DNA Technologies (IDT) and ligated into the *Aat*II and *Eco*RI sites of the above plasmid; shield mutations to the guide were incorporated to avoid continuous cutting of the genome by Cas9. An identical plasmid was constructed containing a silent mutation at A460 (gct → gcc). Transfection of *Pf*3D7 was performed as previously described (48) and parasites selected for with 2.5 mM WR99210.

#### Thermal proteomics profiling (TPP) of MMV1557817

Parasite isolation and parasite protein solubilization were performed as previously described (7). For experiment 1, parasite lysate was separated into technical replicates: four for DMSO control (0 nM) and three for 300 nM of **MMV1557817** treatment. For experiment 2, parasite lysate was divided into four technical replicates of DMSO control (0 nM) and 1200 nM of **MMV1557817** treatment. The samples were then incubated at room temperature for 3 min before being thermally challenged by heating at 60°C for 5 min. The denatured protein fraction was then removed via ultracentrifugation at 100,000 *g* for 20 min at 4°C (Beckman Coulter Optima XE-90 – IVD ultracentrifuge with a 42.2 Ti rotor). The soluble fraction was then processed for proteomics analysis as previously described (7, 49). LC-MS/MS was analysed using data-independent acquisition mode as previously described (49). Raw files were processed using in-house generated *P. falciparum* spectral library using SpectronautTM 13.0 as previously described (49). The relative abundance of identified proteins were calculated as fold change of drug-treated conditions relative to the 0 nM control for each experiment (only for proteins with intensities greater than 1×10^5^ and with a minimum peptide count of 2). Significant proteins determined by Welch’s *t*-test (p-value <0.05) and fold change >1.2, at multiple concentrations of **MMV1557817** were considered as stabilized proteins and were plotted using paired volcano plots.

#### Metabolomics analysis of MMV1557817, MIPS2673 and 3 compared to DMSO control

*P. falciparum* (3D7) cultures at 6% parasitaemia and 2% haematocrit were subject to double sorbitol synchronisation 14 hours apart, followed by further incubation for 28-42 h to achieve the desired trophozoite stage (28 hours post infection). Infected RBCs (2×10^8^) were treated with 10× the IC_50_ of MMV1557817 (320 nM), MIPS2673 (1 µM), **3** (4.53 µM) for 1 h, after which metabolites were extracted, analysed and processed as previously described (7). Principal-component analysis (PCA) and hierarchical clustering algorithms were run also in Metaboanalyst (50). Metabolomics data are presented as relative abundance values from 4-7 biological replicates. Differences were determined using Welch’s *t*-test where significant interactions were observed. Significance was determined at p values < 0.05. The metabolomics data for **3**, MIPS-2673 and drug-free controls were reported previously (7, 51).

#### Blue Native PAGE and Western blotting

Erythrocytes infected with parental Dd2 or **MMV1557817** resistant parasites were grown to trophozoite stage and 60 mL of culture with greater than 5% parasitaemia was lysed with 0.1% (w/v) saponin in PBS and washed 3 times to remove haemoglobin. Following centrifugation, the parasite pellet was solubilized with 1% (v/v) Triton X-100, then incubated on ice for 30 min. Insoluble material was pelleted (14 000 *g* for 30 min at 4°C). The protein concentration of the supernatant was determined by Bradford assay (BioRad). Next, the supernatants were electrophoresed on NativePAGE Novex 3–12% Bis-Tris protein gels as per manufacturer’s instructions (Invitrogen). Briefly, proteins were separated at 150-200 V at 4°C until the dye front ran off the gel. Proteins were transferred to methanol-activated PVDF membrane using a wet-transfer with a constant current of 300 mAmp for 90 min. Membranes were incubated with 8% acetic acid in water for 20 min at room temperature before being rinsed with MilliQ water and dried overnight. Once dried, the membrane was washed in methanol and Western blotting was performed. Blots were blocked in 3% (w/v) bovine serum albumin (BSA) in PBS before being probed with rabbit anti-M17 (1:1000) as previously described (7). Horseradish peroxidase-conjugated secondary antibodies were used (1:10,000; Thermo Scientific) and protein bands were detected using the Clarity ECl Western blotting substrate (BioRad) and imaged on a BioRad ChemiDoc^TM^ Imaging System

#### Visualization of aminopeptidase activity in live cells using fluorogenic peptide substrates

Aminopeptidase activity within live Dd2 parasites was determined using a fluorogenic H-Leu-NHMec peptide (Sigma) as described previously (52). Briefly, sorbitol synchronized parasites at trophozoite stage were pelleted, washed twice with complete RPMI media and incubated with 10 µM H-Leu-NHMec or N-terminally blocked Z-Arg-ArgMec. After 10 min incubation at 37°C, 10 µl of treated parasites were spotted onto a glass slide, covered with a coverslip, and imaged using a DAPI filter set on a Nikon Eclipse Ti2 microscope at 100 x magnification under oil immersion. To inhibit *Pf*A-M1 activity, washed parasites were treated with MIPS2673 (3.2 µM; 10 x IC_50_ (51)) for 20 min prior to substrate addition. The relative fluorescence detected within a parasite cytosol was calculated using ImageJ (NIH, version 1.53c) and was expressed relative to surface area. Images of at least 20 individual parasites across 2 independent experiments were taken per treatment under identical conditions. Inhibitors and substrates were diluted in DMSO; total DMSO did not exceed 0.5% of final volume.

#### Parasite haemoglobin fractionation assay

Haemoglobin was fractionated as previously described with some alterations (53). Double sorbitol synchronized Dd2 or **MMV1557817** resistant parasites at mid-late trophozoite stage were harvested with 0.1% saponin in 1×PBS containing 1× protease inhibitors at 4°C for 10 min before undergoing 3× washes in 1×PBS containing 1× protease inhibitors. Pellets were sonicated in 50 µL of water for 5 min after which 50 µL of 0.2 M HEPES (pH 7.5) was added and the mixture was centrifuged at 1500 *g* for 20 min. The supernatant was processed further by addition of 50 µL 4% SDS (w/v) and a further 5 min sonication. The samples were then incubated at 95°C for 5 min before addition of 50 µL of 0.3 M NaCl and 50 µL 25% (v/v) pyridine in 0.2 M HEPES (pH 7.5) and vortexing; this sample contained the haemoglobin fraction. The pellet was resuspended in 50 µL water and 50 µL 4% SDS (w/v) and sonicated for 5 min before being incubated at 95°C for 5 min. To this, 50 µL of 0.2 HEPES (pH 7.5), 50 µL 0.3 M NaCl and 50 µL 25% pyridine (v/v) was added and the sample vortexed and then centrifuged at 1500 *g* for 10 min; the resulting supernatant contained the haem fraction. The remaining pellet was solubilized in 50 µL water and 50 µL 0.3 M NaOH by sonication for 5 min and incubation at 95°C for 5 min. Finally, 50 µL of 0.2 M HEPES (pH 7.5), 0.3 M HCl and 50 µL 25% pyridine (v/v) was added and samples were vortexed; this sample contained the hemozoin fraction. The absorbance of samples was measured at 405 nm on a Perkin Elmer Ensight Plate Reader.

## RESULTS

### MMV1557817 is active against aminopeptidases from multiple *Plasmodium* spp

We confirmed inhibitory constants of **MMV1557817** to be in the nanomolar range for all six relevant aminopeptidases from key *Plasmodium* clinical (*P. falciparum, P. vivax)* and animal (*P. berghei*) models (*Pf*A-M1, *Pf*A-M17*, Pv*-M1*, Pv*-M17*, Pb*-M1 and *Pb*-M17; Fig 1A), before the binding mode of the compound was investigated. Data was obtained from co-crystallographic structures of the *P. falciparum* enzymes (*Pf*A-M1, *Pf*A-M17) and was considered representative of the homologous proteins due to the conservation of active site residues and architecture (Supplementary Table 1; (16)). In M1, the 3,4,5-trifluoro bi-phenyl moiety occupied the S1 subsite with similar interactions as reported previously, including a carbonyl-π with the main chain oxygen of Glu319, edge-face π-stacking with Tyr575 and Met1034 packing against the fluorinated ring (6) (Fig 1B). The interaction with Met1034 involves movement of the sidechain compared to its unbound position. The S1’ moiety of **MMV1557817** is a 3,3-dimethylbutanamide that occupies a similar position in the S1’ subsite, making limited contacts with residues lining the pocket. For the M17 enzymes, the large 3,4,5-trifluoro bi-phenyl moiety forms interactions with residues lining both the sides and top of the S1 subsite (Fig 1C). The methionine side chain situated at the top of the S1 subsite extended away from **MMV1557817**, likely displaced by the bound compound. Variations in this Met sidechain position are likely dependent on the degree of rotation adopted by the biphenyl group. The 3,3-dimethylbutanamide group extends into the S1’ subsite and shows little flexibility.

**Figure 1.**
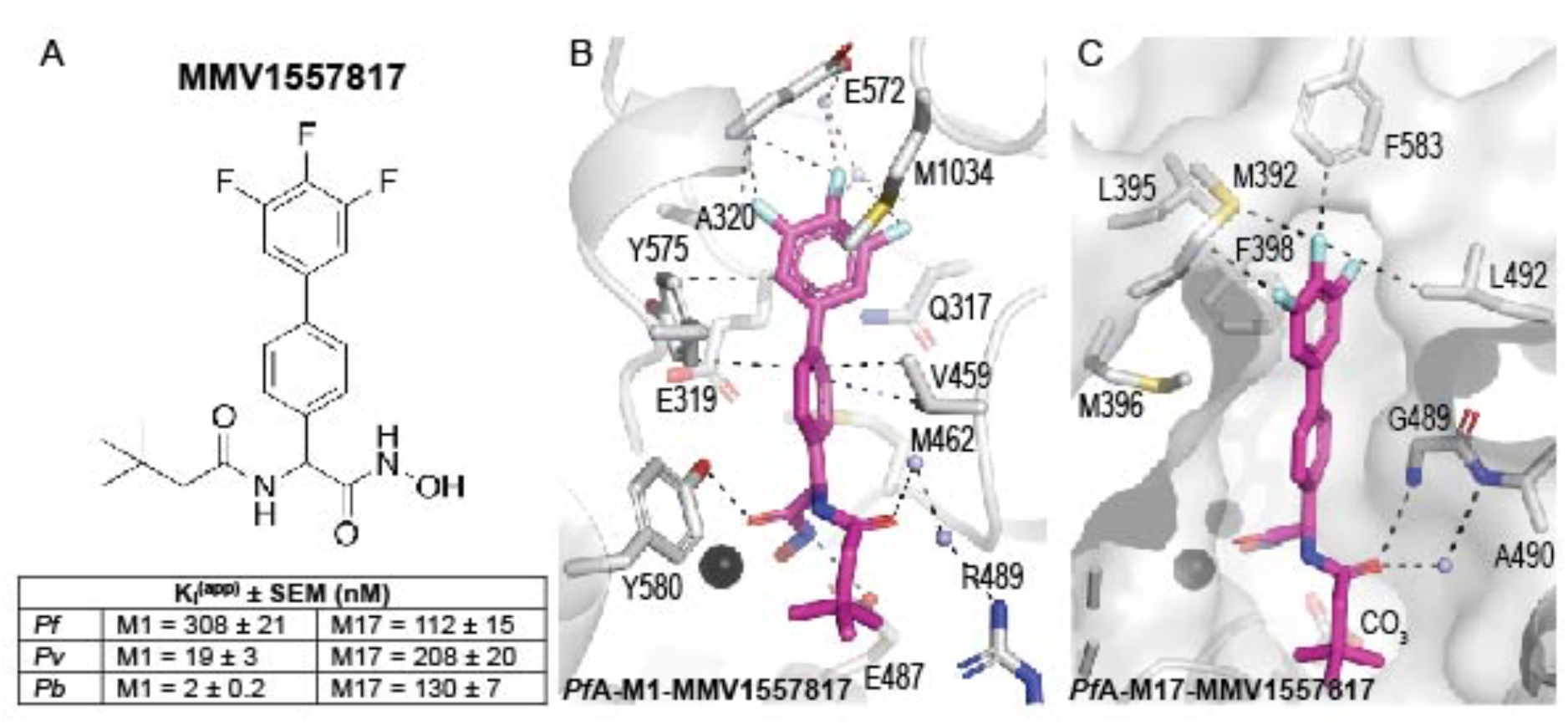
MMV1557817 is a potent dual inhibitor of *Plasmodium* M1 and M17 aminopeptidases. (A) Chemical structure of **MMV1557817**. Inhibition constants (*K*_iapp_, in nM) for **MMV1557817** toward recombinant, purified *Pf*A-*Pv*- and *Pb*-M1 and *Pf*A-*Pv*- and *Pb*-M17 are shown underneath panels with the standard error of mean (SEM) indicated. (B, C) Binding mode of **MMV1557817** to *Pf*A-M1 (B) and *Pf*A-M17 (C). Interactions between **MMV1557817** (magenta sticks) and protein residues (grey) shown as black dashed lines. Residues of interest are indicated.

### MMV1557817 demonstrates potent activity against asexual and sexual stage parasites

Preliminary characterisation of **MMV1557817** against *P. falciparum* asexual *Pf*3D7 and Dd2 parasites was performed previously using an image based screening assay (54), and the IC_50_ found to be in the low nanomolar range (11). To determine the effect of **MMV1557817** on parasite growth we investigated the stage at which growth was affected after treatment. Using the IC_50_ determined (Fig 2A; 39 nM, 31.8-46.9 CI), *Pf*3D7 cultures at 0 – 4 h old were treated with 5× or 10× the IC_50_. Both treatments resulted in stalling of parasite growth at the ring stage, while parasites treated with the 10× concentration of DMSO continued through the cycle before reinvading into the following cycle approx. 48 h after treatment began (Fig 2B).

**Figure 2.**
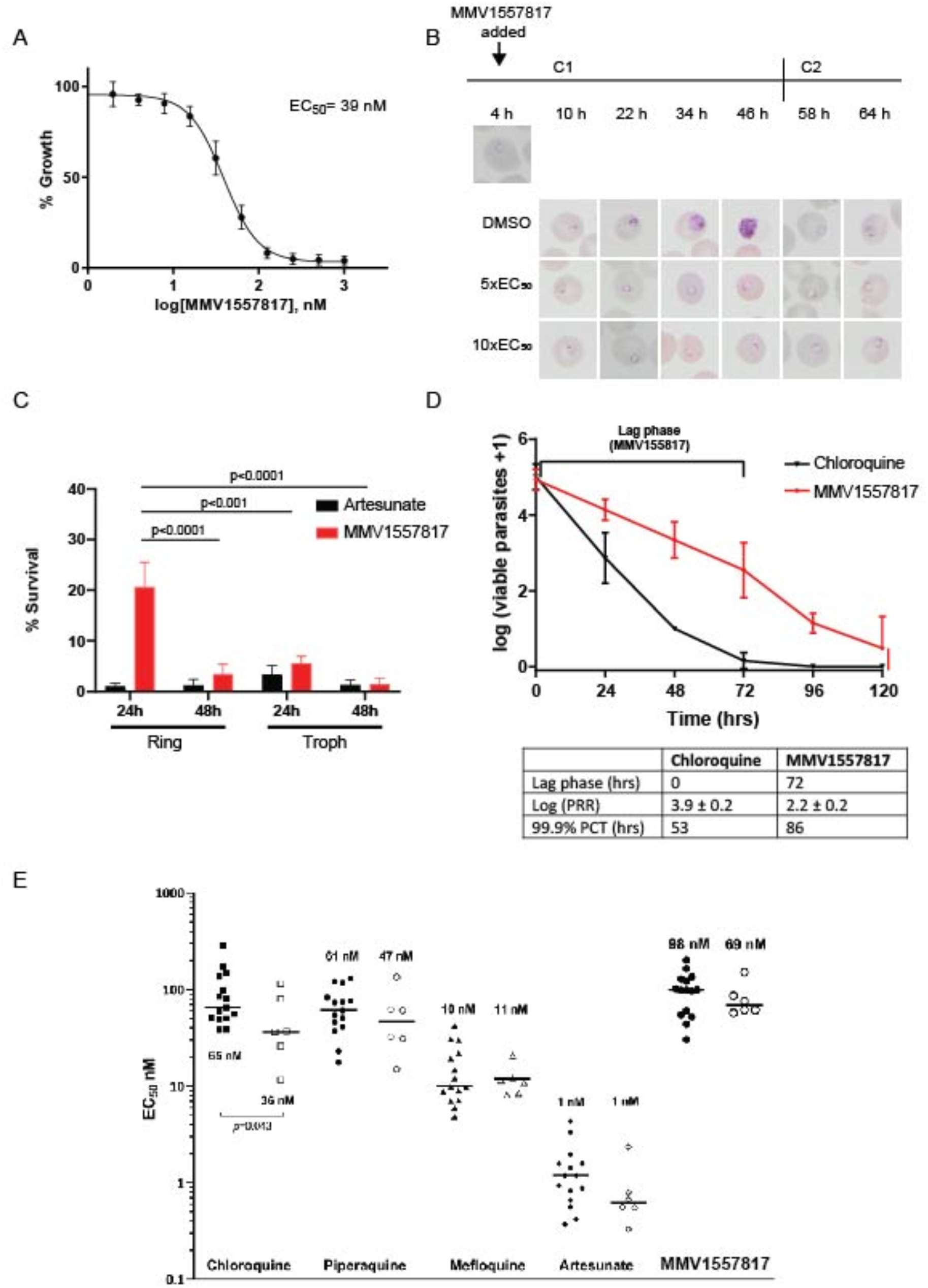
Activity of MMV1557817 against *P. falciparum* and *ex vivo P. vivax*. A) Killing action of **MMV1557817** determined by a standard 72 h ring killing assay. Plotted is the mean ± standard error of the mean (n=3 performed in triplicate). (B) Representative Giemsa-stained Pf3D7 parasites treated with **MMV1557817** over two cycles (C1, cycle 1; C2, cycle 2) beginning at 4 hours post infection (hpi) with 5× or 10× the IC_50_ or DMSO at the concentration present in the 10× IC_50_ treatment. Treated parasites did not progress past ring stage. (C) Assessment of **MMV1557817** activity on ring and trophozoite stage *Pf*3D7. Parasites were incubated in 10 x IC_50_ **MMV1557817** for either 24 or 48 hr before the compound was washed off and parasites allowed to grow for a further 48 hr. Survival was determined via Sybr Green I assay and compared to vehicle (DMSO)-treated controls. Shown is the mean ± standard deviation (n=4). Statistical significance was determined using a one-way ANOVA. (D) *Pf*3D7 viability time course profiles for **MMV1557817** compared to chloroquine. The lag phase is indicated on the graph and summarised in the table below, along with the log of the parasite reduction ratio (PRR) and the 99.9% parasite clearance time (PCT; n≥2 performed in quadruple). (E) Killing action of **MMV1557817** and current antimalarial compounds against *ex vivo P. falciparum* and *P. vivax* obtained from clinical isolates. No significant differences in IC_50_ values were observed for these compounds between species, with the exception of chloroquine.

Wash-out experiments were performed to further analyse asexual killing, with artesunate serving as a positive control (IC_50_ 4 nM, 0.5-6.5 CI). Treatment with **MMV1557817** for 48 h, starting at ring stage, was significantly more effective at killing parasites than a 24 h exposure. When treatment was initiated at trophozoite stage, 24 h exposure to **MMV1557817** was significantly more effective than a similar exposure at ring stage (Fig 2C). With the exception of the 24 h treatment of rings, there was no significant difference between parasite survival after treatment with 10× IC_50_ of **MMV1557817** or artesunate for the different time periods (Fig 2C).

Next, the parasite reduction ratio (PRR) and parasite clearance time (PCT) of **MMV1557817** were assessed *in vitro*, as both represent important mode-of-action parameters to determine the likelihood of parasite recrudescence and drug resistance *in vivo*. **MMV1557817** showed a lag phase of 72 hours before reaching its optimal killing rate. After this lag phase, the PRR over one cycle was 2.2 ± 0.2 and the 99.9% PCT was 86 hours (Fig 2D). The antimalarial chloroquine was used as a positive control, with a PRR of 3.9 ± 0.2 and the 99.9% PCT of 53 hours, similar to what has been described previously (35).

The ability of **MMV1557817** to kill the sexual stages of *P. falciparum* was also determined using image-based assays. The IC_50_ values for early (stage I-III; n=3), late (stage IV-V; n=3) and mature (stage V; n=2) gametocytes were determined to be 99 nM, 309 nM, and 1474 nM, respectively (Table 1). While there is a reduction in the activity against the sexual stages compared to the asexual stages, **MMV1557817** is nevertheless still effective against early and late-stage gametocytes at sub-micromolar concentrations.

**Table 1:** Activity of MMV1557817 against *P. falciparum* gametocytes (IC_50_ values (nM); mean ± SD)

| Compound | Early-stage Gametocytes | Late-stage Gametocytes | Mature-stage Gametocytes |
| --- | --- | --- | --- |
| **MMV1557817** | **99 ± 2.0** | **309 ± 1.0** | **1474 ± 304** |
| artesunate | 16 ± 5.0 | 26 ± 2.0 | 51 ± 23 |
| pyrimethamine | N/A at 40µM | N/A at 40µM | N/A at 40µM |
| chloroquine | 157 ± 3.5 | 99% at 40µM | 94% at 40µM |
| pyronaridine | 67 ± 7.0 | 2280 ± 679 | 1254 ± 120 |
| puromycin | 163 ± 18 | 94 ± 5.0 | 73 ± 24 |
| DHA | 16 ± 2.0 | 35 ± 12 | 103 ± 30 |
| Methylene Blue | 110 ± 4.0 | 52 ± 2.0 | 63 ± 26 |

### MMV1557817 retains nanomolar range activity against drug resistant and clinical isolates of *P. falciparum* and *P. vivax*

To determine the efficacy of **MMV1557817** against known drug resistance mechanisms, activity was tested against a panel of drug resistant field isolates using the previously described [^3^H]hypoxanthine incorporation assay (34) and compared to the chloroquine (CQ) sensitive field isolate NF54 (IC_50_ = 22 nM; Table 2). The strains that were tested include K1 (CQ, pyrimethamine and sulfadoxine resistant), 7G8 (CQ resistant), TM90C2B (atovaquone resistant), Cam3.1 (artemisinin resistant) and Dd2 (CQ resistant; previously tested (11)). No significant shift in **MMV1557817** IC_50_ was observed when compared to the NF54 strain, indicating that the compound is not susceptible to any known resistance mechanism tested here (Table 2). Importantly, no loss of efficacy was demonstrated for the artemisinin resistant Cam3.1 strain, suggesting that the drug would be effective against artemisinin resistant parasites. We additionally investigated the fractional inhibitory concentrations (FIC) of artemisinin and **MMV1557817** used in combination on Dd2 parasites to determine any potential interactions between the two compounds. The ΣFIC of each ratio combination found the compounds to function in an additive manner, whereby each compound retained their activity in the presence of the other (Supplementary Fig 1 and Supplementary Table 2). **MMV1557817** was further tested against laboratory selected Dd2 parasites that harboured additional resistance mutations to current novel antimalarials including DDD107498 (protein synthesis inhibitor (55)), MMV390048 (phosphoatidylinositol 4-kinase inhibitor (56)), DSM265 (dihydroorotate dehydrogenase inhibitor (57)), GNF156 (*Pf*carl (58)) and ELQ300 (cytochrome bc1 complex inhibitor (59)), and again showed no IC_50_ shift compared to the parent Dd2 line indicating that **MMV1557817** is not targeting these mechanisms.

**Table 2:**
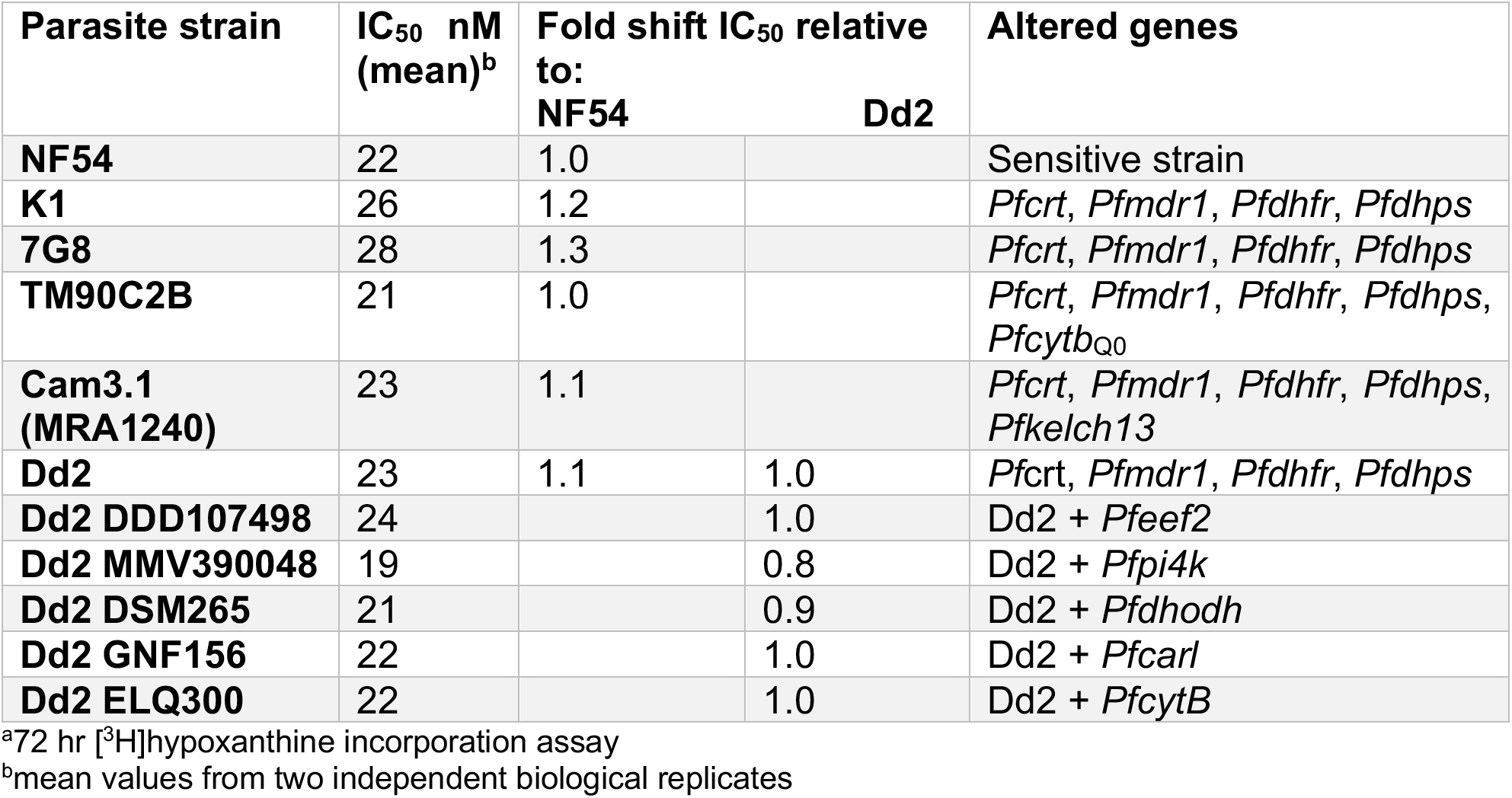
MMV1557817 effectiveness against drug resistant *P. falciparum* strains^a^.

To assess the efficacy of **MMV1557817** against clinical isolates, *ex vivo* susceptibility was performed on isolates from 23 patients presenting to malaria clinics in Timika (Indonesia) with single-species infections of either *P. falciparum* (n=15) or *P. vivax* (n=8) between January and April 2016. Susceptibility profiles in the same isolates were also determined for standard anti-malarials CQ, piperaquine, mefloquine and artesunate. Adequate growth after harvesting was achieved for all (15/15) of *P. falciparum* isolates and 75% (6/8) of *P. vivax* isolates. Baseline characteristics of the isolates are presented in Supplementary Table 3. Drug susceptibility did not differ significantly between species for the anti-malarial drugs piperaquine, mefloquine and artesunate, as well as **MMV1557817** (median IC_50_: 98.1 nM for *P. falciparum* versus 68.6 nM for *P. vivax*; p=0.533; Fig 2E and Supplementary Table 4). However, significantly greater activity was observed against *P. vivax* than *P. falciparum* for CQ (median IC_50_: 64.8 nM for *P. falciparum* versus 36.4 nM for *P. vivax*; p=0.043). The nanomolar efficacy of **MMV1557817** against *P. vivax* confirms cross species killing consistent with previously generated *K*_i_ values against *P. falciparum* and *P. vivax* recombinant M1 and M17 proteins (11).

### MMV1557817 shows good selectivity, safety, ADME and exposure properties and is active *in vivo* against the rodent malaria species *Plasmodium berghei*

Previously, the selectivity of **MMV1557817** was assessed against matrix metalloproteinase (MMP) 2, 7, 8, 9 and 13, as well as the human M1 aminopeptidase insulin regulated aminopeptidase (IRAP) and the human aminopeptidase N (APN or CD13) (11). Collectively, these studies identified that **MMV1557817** shows limited off-target inhibitory affects against the MMPs and the IRAP (tested up to 200 µM). For the human APN, some cross-reactivity with a *K*_i_ of 0.3 µM was observed. Given this cross-reactivity with human APN, we further assessed the efficacy of **MMV1557817** toward other human M1 aminopeptidases of current or future therapeutic interest (60). Here we tested human leukotriene A4 hydrolase (LTA4H) and the endoplasmic reticulum aminopeptidases 1 and 2 (ERAP1/2). These enzymes were purchased commercially and showed limited activity in our *in vitro* assay and as such, an appropriate 12-point dose range in triplicate was not viable and a *K*_i_ calculation could not be completed. Thus, the percent of enzyme activity was compared in the presence of different concentrations of **MMV1557817** or artesunate (Table 3). The results showed that at 1.25 µM compound, **MMV1557817** resulted in 18% inhibition of LTA4H but had no activity toward either ERAP. At the highest concentration tested (1 mM), there was near complete inhibition of all three human homologs. Artesunate showed minimal inhibition on the activities of the human enzymes but did show moderate, dose-dependent inhibition of the *Plasmodium* M1 enzymes. Our results suggest that there is some cross-reactivity to human homologs, however, the selectivity index between human and *Plasmodium* target remains high (50–100).

**Table 3:** Inhibition of aminopeptidase activity with increasing concentrations of MMV1557817 or artesunate.

| [Compound]<br>(µM) | *Pf*-M1 | *Pv*-M1 | *Pb*-M1 | LTA4H | ERAP1 | ERAP2 |
| --- | --- | --- | --- | --- | --- | --- |
| **MMV1557817** |  |  |  |  |  |  |
| 1.25 | 83 % | 96 % | 94 % | 18 % | 0 % | 7 % |
| 10 | 100 % | 100 % | 100 % | 16 % | 0 % | 0 % |
| 500 | - | - | - | 71 % | 75 % | 99.1 % |
| 1000 | - | - | - | 87 % | 98 % | 100 % |
| **Artesunate** |  |  |  |  |  |  |
| 1.25 | 4.8 % | 2.1 % | 0 % | 0 % | 0 % | 0 % |
| 10 | 11.9 % | 23.6 % | 12.8 % | 12.8 % | 0 % | 0 % |
| 500 | 16.8 % | 74.0 % | 81.3 % | 20.2 % | 0 % | 0 % |
| 1000 | 43.2 % | 68.2 % | 78.2 % | 36 % | 0 % | 0 % |

*In vitro* pharmacology safety assays conducted using the EuroFins CEREP panel were used to assess potential off-target activity within DNA repair pathways (namely the human histidine deacetylases HDAC1, 2, 5, 7, 8, 9, 10) and protease pathways (human matrix metalloproteases MMP-2, −3, −7, −8, −9, −14) (Supplementary Table 5). Overall, the results only identified MMP-8 as being just above the significance threshold for inhibition by **MMV1557817** at 10 µM (52.5% inhibition), with no significant stimulation or inhibition of any of the receptors tested.

Studies to assess physicochemical properties, plasma stability and metabolic stability were performed with **MMV1557817** prior to assessing the exposure profile in mice. As shown in Table 4, the compound has a moderate molecular weight and polar surface area, a moderately high Log D_7.4_, and low to moderate aqueous solubility. It was reasonably stable in liver microsomes and cryopreserved hepatocytes and in mouse and human plasma. The compound was highly bound to both plasma proteins (98-99% bound) and in the Albumax medium (84.5% bound) used for *in vitro* activity assessment. Correcting for the binding in Albumax medium gives an unbound IC_50_ value of 3.4 nM (based on NF54, Table 2).

**Table 4:** Physicochemical and *in vitro* stability properties of MMV1557817.

| Parameter | MMV1557817 |
| --- | --- |
| MW | 394.39 |
| calc Log D<sub>7.4</sub> | 3.6 |
| Kinetic Solubility (µg/mL) pH 2.0 / 6.5 | 12.5-25 / 12.5-25 |
| *in vitro* microsomal CL<sub>int</sub> (µL/min/mg) H / R / M* | 11 / 29 / 23 |
| *in vitro* hepatocyte CL<sub>int</sub> (µL/min/10<sup>6</sup> cells) H / R* | 2 / 4 |
| Plasma stability (% remaining, 4 h, 37°C) H / M* | 97 / 93 |
| Plasma protein binding (%) H / R / M* | 98.8 / 97.8 / 97.7 |
| Albumax media binding (%) | 84.5 |

H: human, R: rat, M: mouse

Studies with human and rat hepatocytes identified three main primary metabolites (Supplementary Fig 2) corresponding to an amide hydrolysis product (*m*/*z* 380, M-15), a hydroxamic reduction product (*m*/*z* 379, M-16), and a glucuronidation product (*m*/*z* 571, M+176). Authentic standards for these metabolites were not available, which precluded an accurate assessment of the relative abundance of each. However, on the basis of peak area alone (and with the assumption of similar response factors for all metabolites), the hydrolysis product and glucuronide appeared to be the predominant products in human hepatocytes, while the hydroxamic reduction product and the glucuronide were more prominent in rat hepatocytes.

Following this, uninfected mice were treated with a single oral dose of **MMV1557817** at 50 mg/kg administered in a suspension formulation and the plasma concentration versus time profiles as well as plasma exposure parameters were determined (Fig 3A). Unbound concentrations were calculated using the measured free fraction in mouse plasma (Table 4). The half-life was found to be 4.3 h, with the overall results indicating that daily administration at this dose level would be expected to maintain unbound concentrations above the unbound IC_50_ (3.4 nM based on NF54, Table 2) for approximately 14 h.

**Figure 3.**
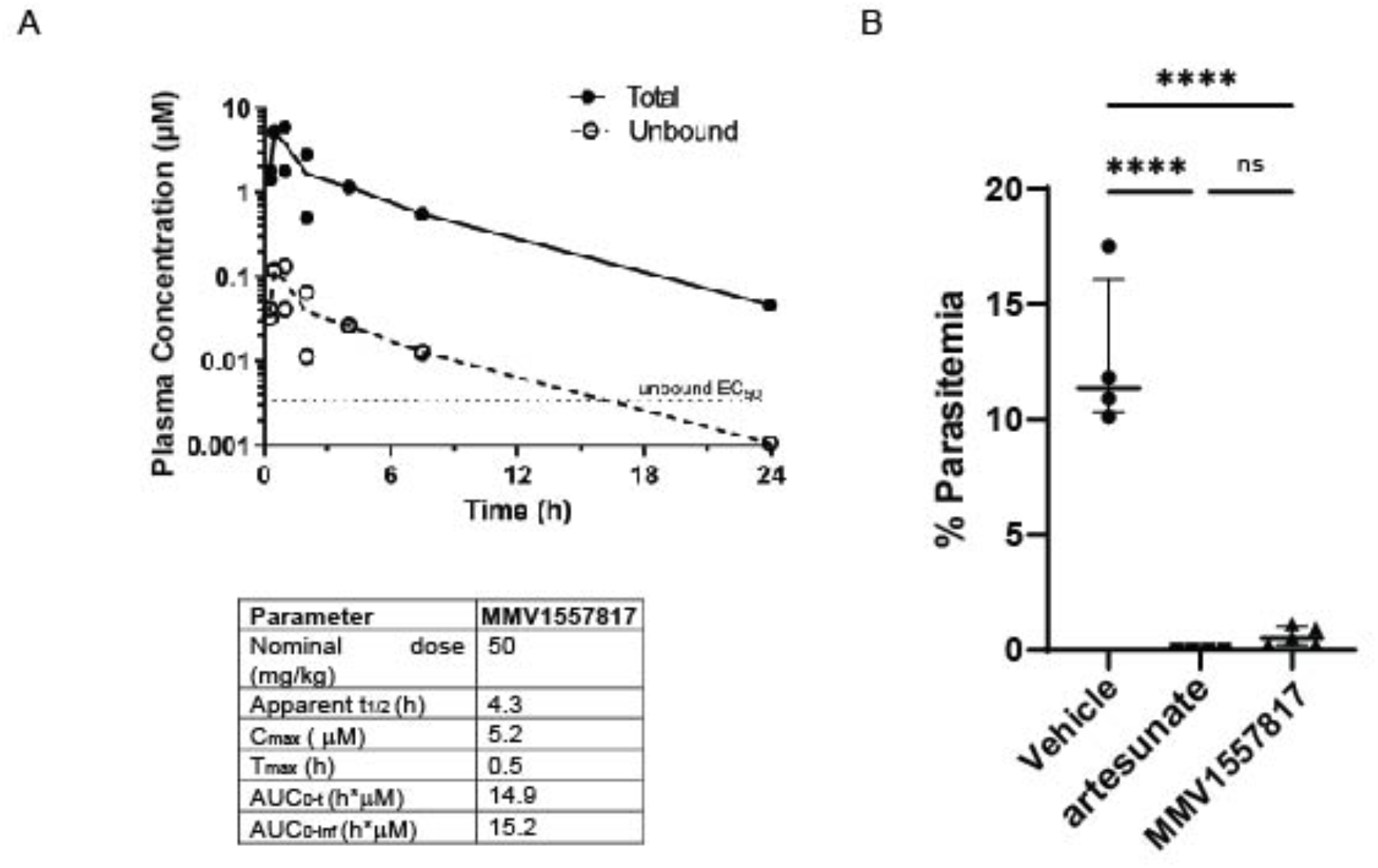
Bioavailability and activity of MMV1557817 in murine models. (A) Upper panel: Plasma concentration versus time profiles for **MMV1557817** following oral dosing (50 mg/kg) to non-infected male Swiss outbred mice. Concentrations are the total measured concentrations and calculated unbound concentrations (obtained by multiplying the total concentration by the fraction unbound in mouse plasma). The unbound *P. falciparum* IC_50_ value for **MMV1557817** is shown for comparison. Lower panel: Plasma exposure parameters in Swiss outbred mice following oral administration. (B) Peter’s test performed on Balb/c mice infected with *P. berghei* ANKA parasites and treated with 50 mg/kg **MMV1557817** or 30 mg/kg artesunate on days 0,1,2,3 post-infection and parasite clearance determined on day 4. **MMV1557817** was found to clear 95.4% of infection. Significance was determined using an unpaired t-test; n≥4.

Given the good selectively and reasonable PK properties of **MMV1557817** and after confirming the effectiveness of the compound against recombinant *Pb*-M1 and *Pb*-M17 (Fig 1A), the efficacy of **MMV1557817** was also tested in an *in vivo* model of malaria infection using the *P. berghei* ANKA rodent malaria four-day suppressive test (46). Accordingly, 50 mg/kg **MMV1557817** or 30 mg/kg artesunate (positive control) was given by oral gavage (same formulation as used for the mouse exposure study) on day 0,1, 2 and 3 post-infection, with survival calculated on day 4. **MMV1557817** showed excellent efficacy, with mice treated with this compound exhibiting a 95.4% parasite reduction when compared to the vehicle control, with no significant difference seen between **MMV1557817** treatment and the artesunate control (Fig 3B).

### Thermal proteomics profiling (TPP) and metabolomics confirms MMV1557817 targets both *Pf*A-M1 and *Pf*A-M17 aminopeptidases

**MMV1557817** shows aminopeptidase inhibition of both *Pf*A-M1 and *Pf*A-M17 in recombinant systems, however, recently published resistance selection studies only identified *Pf*A-M17 as a target (61). To elucidate any other potential targets of **MMV1557817** within parasites, thermal proteomics profiling (TPP) was utilized. TPP is based on the principle that, when heated, parasite proteins will denature and can be removed by ultracentrifugation (100,000 *g*) due to their insolubility. In contrast, any protein bound to **MMV1557817** will exhibit enhanced thermal stability, protecting it from the thermal challenge and resulting in an increased concentration of protein in the soluble fraction, which can be identified by LC-MS-based proteomics analysis. Two TPP experiments were performed with 3-4 technical replicates and a thermal challenge of a single temperature at 60°C with low (300 nM (10 x IC_50_), experiment 1), and high (1200 nM (40 x IC_50_), experiment 2) compound concentrations. Common proteins (1405) between experiment 1 (1586) and experiment 2 (1853) identified by LC-MS/MS were analysed and only two proteins had altered thermal stability profiles (fold change >1.2 and p-value <0.05) across the two concentrations of **MMV1557817** compared to the DMSO vehicle treated lysates (0 nM; Fig 4A). These proteins were *Pf*A-M1 (PF3D7_1311800) and *Pf*A-M17 (Pf3D7_1446200). *Pf*A-M1 demonstrated a 1.36-fold stabilisation following incubation with 300 nM **MMV1557817** and a 4.33-fold stabilisation with 1200 nM compared to 0 nM. *Pf*A-M17 demonstrated a significant 1.28-fold stabilisation with 300 nM **MMV1557817** treatment, and 1.26-fold stabilisation with 1200 nM compared to 0 nM (Fig 4B). This independent and unbiased drug target identification approach identified *Pf*A-M1 and *Pf*A-M17 as the only protein targets of **MMV1557817**.

**Figure 4.**
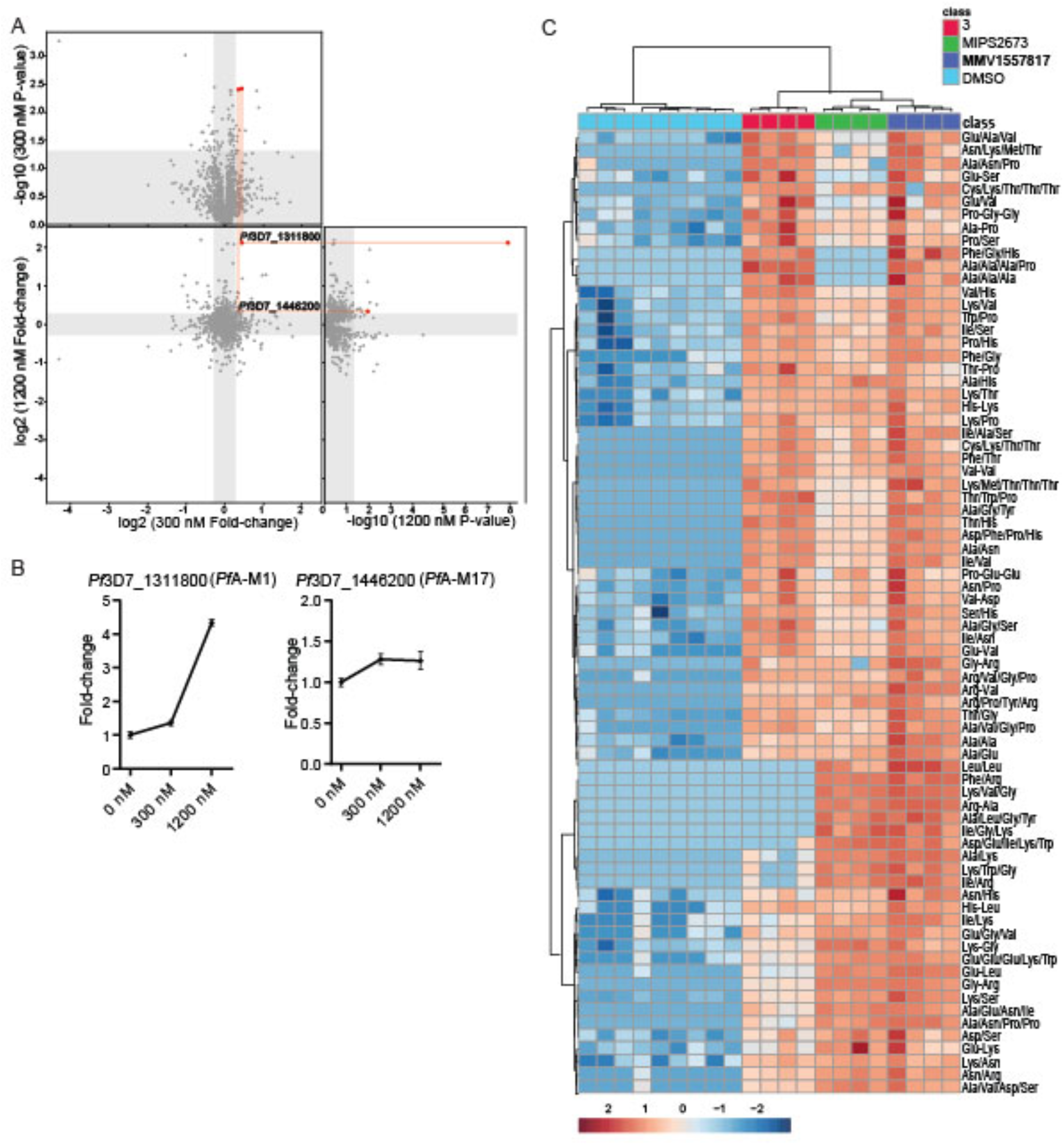
Thermal proteome profiling of MMV1557817 confirms PfA-M1 and PfA-M17 as compound targets. (A) Paired volcano plot of all proteins detected. The outside panels show the log_2_ fold change vs –log_10_ p-value of proteins following treatment with 300 nM or 1200 nM of **MMV1557817**, relative to the 0 µM negative control, following a 60°C thermal challenge. The thermal stability of both PfA-M1 (Pf3D7_1311808) and PfA-M17 (Pf3D7_1446200) were altered at both concentrations with a *p*-value <0.05 as determined by Welch’s *t*-test, with increasing stability in increasing concentrations. *Pf*A-M1 and *Pf*A-M17 are highlighted with red lines and dots. No other proteins were significantly stabilised by both drug concentrations. (B) Protein intensity of *Pf*3D7_1311808 (*Pf*A-M1) and *Pf*3D7_1446200 (*Pf*A-M17) in increasing concentration of **MMV1557817** and following 60°C thermal challenge. Value represents the mean of two biological replicates performed with ≥3 technical replicates ± standard deviation. (C) Hierarchical clustering of the 73 significantly perturbed peptides (*p*-value <0.05) following treatment with **MMV1557817**, MIPS2673 (*Pf*A-M1 inhibitor (51)), **3** (*Pf*A-M17 inhibitor (7)), and DMSO control. Vertical clustering displays similarities between samples, while horizontal clusters reveal the relative abundances of the 73 peptides from 4-7 biological replicates. The colour scale bar represents log2 (mean-centred and divided by the standard deviation of each variable) intensity values. Peptides with hyphen (-) notations indicate confirmed sequence by MS/MS. Peptides with slash (/) notation indicate putative amino acid composition (accurate mass), without confirmed sequence order.

To further confirm that **MMV1557817** targets *Pf*A-M1 and *Pf*A-M17, we compared metabolomics profiles of *Pf*3D7 parasites treated with 10× IC_50_ of **MMV1557817**, MIPS2673 (*Pf*A-M1 inhibitor (51)), **3** (*Pf*A-M17 inhibitor (7)) and DMSO control (Supplementary Fig 3). Principal component analysis and heatmap analysis of relative abundances of putative metabolites dysregulated following treatment with the inhibitors demonstrated that the most prominent metabolomic signature shared between them was a series of peptides that were increased (Fig 4C; Supplementary Fig 3). Further detailed analysis of dysregulated peptides shared among the inhibitors demonstrates that **MMV1557817** increases the levels of peptides that also increase in abundance following specific inhibition *Pf*A-M1 and *Pf*A-M17 (Fig 4C), consistent with additive inhibition of both aminopeptidases. We have previously shown that majority of these peptides could be mapped to Hb sequences (7, 51).

### A single nucleotide polymorphism (SNP) in *Pf*A-M17 identified in MMV1557817 resistant parasites impacts enzyme hexamerisation

Whilst the targets of **MMV1557817** were confirmed using the above methods, it is also important to assess the ability of parasites to become resistant to the compound and the potential impact this has on other antimalarial drugs. Recent *in vitro* resistance selection and whole genome analysis undertaken to determine if resistance against **MMV1557817** could be selected in Dd2 parasites identified a A460S mutation in one of its intended targets, *Pf*A-M17, as well as a N846I mutation in an AP-3 β subunit (PF3D7_0613500) and a M317I mutation in a non-essential conserved *Plasmodium* protein (PF3D7_1144400) (61). These three mutations were present in all clones obtained from multiple flasks, with parasites only displaying modest IC_50_ shifts of between 1.5 – 2.9x, suggesting a low level of resistance (61). To confirm whether the A460S mutation was responsible for resistance, CRISPR/Cas9 technology was utilised to attempt to introduce this mutation into *Pf*A-M17 in *Pf*3D7 wildtype parasites using a previously described donor plasmid method (62). However, after 4 weeks, no mutants could be generated despite a silent mutation being incorporated into *Pf*A-M17 within this time (gct to gcc; Supplementary Fig 4) using the same guide RNA. This suggests that the A460S mutation may not be viable in parasites without the other compensatory mutations found in the *in vitro* resistance selection studies or that parasite survival is limited by the apparent fitness cost seen in **MMV1557817** resistant parasites harbouring this mutation (Supplementary Fig 5).

To assess the effect of the A460S SNP on *Pf*A-M17 aminopeptidase activity, we introduced the same mutation into our recombinant gene expression construct and produced the protein *Pf*A-M17(A460S). The mutant protein was purified as per the same protocol used to generate wildtype protein, but it exhibited an altered behaviour during the size exclusion chromatography step, suggesting that the mutation may have affected the ability of the protein to form the biologically active hexamer (16). Attempts to induce hexamer formation via metal supplementation failed to shift the oligomeric equilibrium toward the functional complex as readily as the wildtype (Fig 5A). Assessment of enzymatic activity confirmed this result with increased concentrations of *Pf*A-M17(A460S) required for observable aminopeptidase activity (Fig 5B). To assess if the mutant could form a hexamer, we solved the X-ray crystal structure of the recombinant *Pf*A-M17 (A460S) (Supplementary Table 1). The 2.6 Å structure confirmed the protein can form a hexamer in the crystal lattice and suggests the lack of activity in solution may arise from an instability of the hexamer or oligomeric intermediates. Overall, the structure was virtually unchanged from wildtype (0.351 r.m.s.d over 497 C α atoms within the A-chain; Fig 5C) and no gross changes in secondary or tertiary structure were observed. Further, a similar domain arrangement and no large regions of disorder were observed in electron density that were not already known to be flexible in wildtype structures. The location of residue A460 is in the active site, directly between two highly conserved zinc coordinating residues, D459 and E461. Both D459 and E461 are involved in metal-protein interactions, and E461 shows bidentate zinc coordination when both metal sites are occupied (18). Interestingly, E461 has been shown to be essential for correct hexamerisation and subsequent aminopeptidase activity of *Pf*A-M17 (16). The residue A460 forms a hydrogen bond with the carbonate ion, also present in the active site and required for proteolysis, but the bond is formed with the backbone carbonyl and as such, is maintained when the residue is changed to A460S. The small side-chain A > S change only introduces a hydroxyl group; however, the sidechain does reach into a pocket that is largely hydrophobic. Whether this is enough to destabilise the folding of the pocket and active site, or whether the change interferes with zinc binding (and hence destabilises hexamer formation) remains unclear.

**Figure 5:**
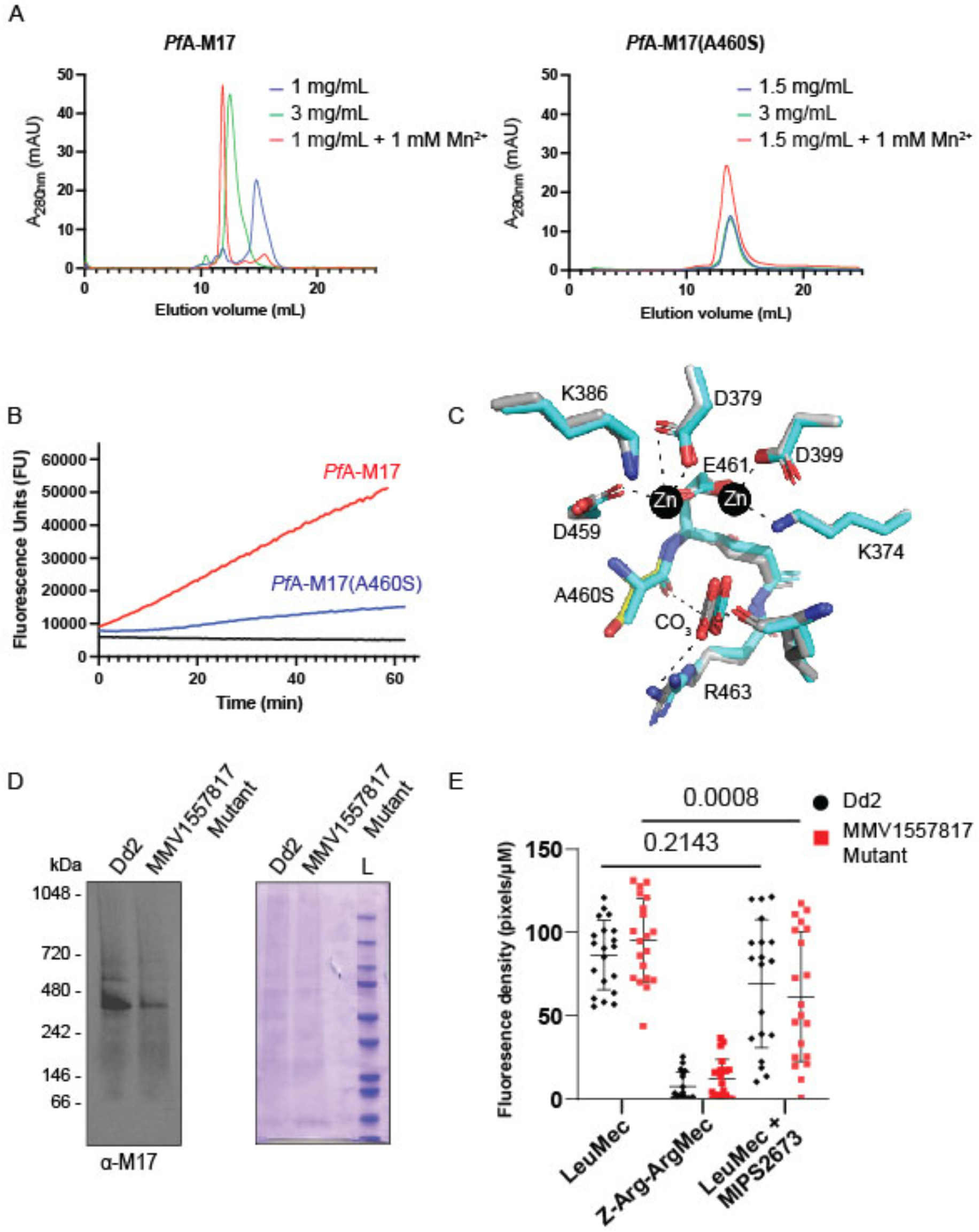
Characterisation of *Pf*A-M17(A460S) (A) Size exclusion chromatography of recombinant wild type *Pf*A-M17 (left) and mutant *Pf*A-M17(A460S) (right) showing that the shift to the functional hexameric species does not occur in the mutant as readily as wild type. Concentration of recombinant proteins analysed is indicated, as is the presence of metal ions that has been shown to shift the oligomeric equilibrium. (B) Activity assays comparing recombinant wild type (red) at 300 nM to A460S mutant (blue) at 10 µM. Aminopeptidase assay is indicated by an increase in fluorescence units (x-axis) over time (y-axis). (C) X-ray crystal structure of the active site of A460S (grey sticks) shows little change compared to wild type (cyan sticks) in zinc-coordinating residues nor interactions with the carbonate ion (labelled), required for activity. The mutation position S460 is shown as a yellow stick. Interactions are shown in dashed lines. (D) Western blot analysis of Blue-Native PAGE performed on trophozoite stage Dd2 or MMV1557817 resistant parasites solubilized in 0.25% Triton X-100 reveals the presence of a *Pf*A-M17 specific species representative of the native homohexamer (left panel). The expected size of the hexamer is 408 kDa and each lane contains 10 µg of protein. Protein loading was confirmed by Coomassie stain (right panel). (E) Quantification of fluorescent density of proteolytic cleavage by *Pf*A-M1 and *Pf*A-M17 in Dd2 or MMV1557817 resistant parasites via live cell imaging with or without *Pf*A-M1 inhibition by MIPS2671. Z-Arg-ArgMec serves as a negative control. Plotted is the mean ± standard deviation and significance determined using a one-way ANOVA Dunnett’s test.

We next compared *Pf*A-M17 in lysates made from wildtype and **MMV1557817** resistant parasites to assess hexamerisation of the protein within parasites. Blue native PAGE confirmed that whilst *Pf*A-M17(A460S) could form hexamers in parasites, reduced amount of hexamers were observed when equal amounts of protein were loaded (Fig 5D). Further, live cell microscopy was employed to investigate the activity of *Pf*A-M17(A460S) in these resistant parasites. The fluorogenic peptide substrate L-Leucine-7-amido-4-methylcoumarin hydrochloride (LeuMec) can be cleaved by both *Pf*A-M1 and *Pf*A-M17 (52), with both Dd2 and **MMV1557817** resistant parasites confirmed to cleave this substrate (Fig 5E and Supplementary Fig 6). The peptide Z-Arg-Arg-7-amido-4-methylcoumarin hydrochloride (Z-Arg-ArgMec) is a N-terminally blocked substrate that is unable to be cleaved and serves as a background fluorescent control. Employing the previously published *Pf*A-M1-specific inhibitor MIPS2673 (51), parasites were treated with 10× the IC_50_ (3.2 µM) for 20 min to determine the level of residual *Pf*A-M17 function. Only **MMV1557817** resistant parasites showed significantly reduced fluorescence with this treatment, suggesting that there is reduced active *Pf*A-M17 present in these parasites, possibly due to the destabilisation of the homohexamer (Fig 5E). This mechanism of resistance is likely to be limited by the essentiality of *Pf*A-M17 (7).

### MMV1557817 resistant parasites show altered susceptibility to aminopeptidase inhibitors and an increase in haemoglobin digestion

To further assess the effect of the SNPs in **MMV1557817** resistant parasites, including the A460S mutation in *Pf*A-M17, we measured the parasite killing of individual aminopeptidase inhibitors against these parasites. The IC_50_ values were determined using the resistant parasites treated with either the *Pf*A-M17 specific inhibitor compound **3** (7) or the *Pf*A-M1 specific inhibitor MIPS2673 (51). Parasites were found to be resistant to the *Pf*A-M17 specific inhibitor, with the IC_50_ showing an increase of 22× when compared to the parental Dd2 parasites (Fig 6A). **MMV1557817** resistant parasites were sensitised to the specific *Pf*A-M1 inhibitor, showing an approximate 3.5× decrease in IC_50_ (Fig 6B). Considering that both enzymes are essential for providing parasites with amino acids from haemoglobin, a decrease in *Pf*A-M17 possibly results in a greater reliance on *Pf*A-M1 for digestion, therefore sensitising parasites to this specific inhibitor.

**Figure 6.**
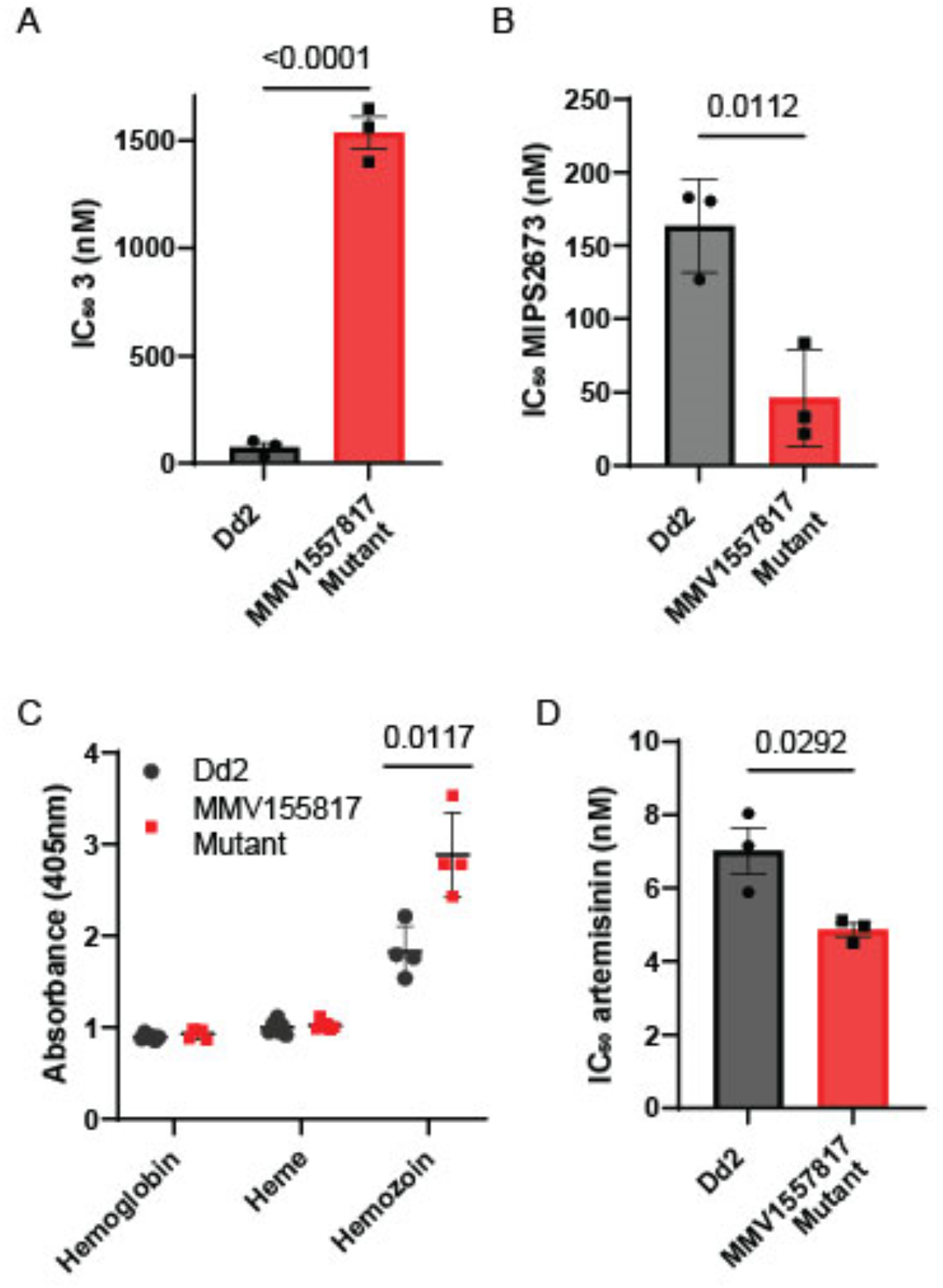
MMV1557817 resistant parasites show altered susceptibility to aminopeptidase inhibitors and artemisinin as well as an increase in haemoglobin digestion. Killing action of an (A) *Pf*A-M17 specific inhibitor (3) or (B) *Pf*A-M1 specific inhibitor (MIPS2673) show altered IC_50_ values compared to parental Dd2 parasites from 3 biological replicates performed in triplicate and plotted as the mean ± standard deviation; significance determined by unpaired *t*-test. (C) Absorbance readings at 405 nm for haemoglobin, haem and hemozoin species from 5.18 x 10^7^ Dd2 or **MMV1557817** resistant parasites at mid-late trophozoite stage. Plotted is the mean ± standard deviation from ≥4 biological replicates; significance determined by Welch’s *t*-test. (D) Killing action of artemisinin showed significantly decreased IC_50_ values for the resistant strain compared to parental Dd2 parasites from 3 biological replicates performed in triplicate and plotted as the mean ± standard deviation; significance determined by unpaired *t*-test.

Considering these changes to aminopeptidase inhibition, as well as the potential impact of the A460S mutation on *Pf*A-M17, we next investigated if there were alterations in haemoglobin digestion in the resistant parasites through haemoglobin fractionation. The quantities of haemoglobin, haem and hemozoin in resistant parasites were compared to the parental Dd2 line at the mid-late trophozoite stage (Fig 6C). While no changes were seen for haemoglobin or haem, there was a significant increase in hemozoin in resistant parasites, with parasites harbouring 1.5× more compared to the Dd2 line (Fig 6C). As the majority of haemoglobin digestion is likely to have already occurred at this later stage of the parasite lifecycle, this increase suggests that there is likely an overall increase in haemoglobin digestion in resistant parasites. Given this, we measured the antiparasitic activity of artemisinin as it requires haemoglobin digestion for drug activation. **MMV1557817** resistant parasites were found to be marginally but significantly sensitized to artemisinin in a 72-hour killing assay (Fig 6D).

## DISCUSSION

As front-line antimalarial treatments continue to be threatened by drug resistance, it is paramount that development of alternative drugs is maintained. Here, we show that M1 and M17 inhibitors are excellent candidates for new antimalarial drugs with novel mechanisms of action. We extensively characterized the dual M1 and M17 aminopeptidase inhibitor, **MMV1557817**, which shows potent, on target activity, using a range of recombinant- and parasite-based assays. The compound showed nanomolar activity toward both *ex vivo P. falciparum* and *P. vivax* parasites, confirming its cross-species activity, as well as a low nanomolar killing range against drug resistant parasites, confirming there is no cross resistance with current known mechanisms. Further, the compound showed selectivity over human analogues and was safely administered in a murine model of malaria, where it was able to clear a *P. berghei* infection. Promisingly, **MMV1557817** shows a low resistance liability, with resistant parasites marginally sensitised to artemisinin, likely due to an increase in haemoglobin digestion. These results suggest that M1 and M17 inhibitors such as **MMV1557817** are worthy of further investigation as potential new antimalarial treatments.

Multiple independent approaches verified that **MMV1557817** targets both *P. falciparum* M1 and M17. Previous studies identified *Pf*A-M17 as a major molecular target for **MMV1557817** (56). We extend this finding using thermal proteomics profiling (TPP), which confirms both *Pf*A-M1 and *Pf*A-M17 are bound by the compound, stabilising these targets at 10 – 40 times their IC_50_ value. Metabolomics analysis of parasites treated with this dual *Pf*A-M1 and *Pf*A-M17 inhibitor found the profile of increased peptides to be a combination of the profiles previously seen in individual enzyme inhibition (7, 51). Taken together, this shows that the primary molecular targets of **MM1557817** are both *Pf*A-M1 and *Pf*A-M17. Resistant parasites generated against **MMV1557817** failed to identify any mutation in *Pf*A-M1, but analysis of the compound’s inhibitory activity may also provide an explanation as to why resistance studies did not isolate a mutation in *Pf*A-M1. In recombinant enzyme assays, **MMV1557817** is two-fold more potent toward *Pf*A-M17 than *Pf*A-M1 (11). The *in vitro* resistance selection in parasites was performed at 90 nM (61), which is similar to the *Pf*A-M17 *K*_i_ (112 nM) but still not as high as the *Pf*A-M1 *K*_i_ (308 nM). In contrast, the TPP was performed at 300 and 1200 nM, both concentrations high enough to bind *Pf*A-M1. We hypothesize that this is the likely explanation for the discrepancy between resistance studies and TPP target identification. Development of **MMV1557817** resistance also only resulted in a very low shift in IC_50_ (1.5 – 2.9x), as well as a significant growth phenotype, suggesting that resistance is not easy to generate, possibly due to the presence of dual targets within the parasite.

The SNP in *Pf*A-M17 identified in **MMV1557817**-resistant parasites alludes to an intriguing mechanism of resistance. Analysis of recombinant *Pf*A-M17 harbouring the A460S mutation suggests that the ability of the protein to form a stable hexamer - essential for aminopeptidase activity - was affected. This also appeared to be the case in resistant parasites, which harboured less *Pf*A-M17 hexamer and showed reduced enzymatic activity. Given that *Pf*A-M17 is essential for parasite survival (7), this mode of resistance is unlikely to lead to full destabilisation of the hexamer as this would likely result in parasite death. Alongside the considerable fitness cost seen in these parasites, it is unlikely that they could persist in an endemic population. Comparison of the binding mode of MMV1557817 and the active site of *Pf*A-M17(A460S) protein suggests that inhibitor binding would not be affected but inhibition could not be assessed due to the lack of activity observed in *in vitro* assays. However, a single A460S mutation in *Pf*A-M17 could also not be reintroduced into *Pf*3D7 parasites, suggesting that the other two SNPs identified may be compensating for a catalytically compromised *Pf*A-M17(A460S) in resistant parasites. Whilst one additional mutation was found in an unknown, non-essential *P. falciparum* protein, the other was identified in the AP-3 β subunit, potentially providing a further layer to **MMV1557817** resistance. This gene is currently only putatively annotated but appears to be essential (20). In mammalian and plant cells, adaptor protein-3 homologs have been shown to be involved in both clathrin-dependent and independent vesicle formation for organelles such as lysosomes, as well as protein cargo sorting in relation to the endoplasmic reticulum and the Golgi apparatus (63–65). The digestive vacuole is analogous to a lysosome, and indeed other adaptor proteins have been identified in the haemoglobin digestion pathway, with AP-3 a notable omission (66). Further investigation into the involvement of the AP-3β subunit in **MMV1557817** resistance may lead to further information on protein trafficking, formation of the digestive vacuole and haemoglobin digestion, where it potentially is responsible for the increase in haemoglobin digestion seen in these parasites.

**MMV1557817** resistance also altered the susceptibility of parasites to individual *Pf*A-M1 and *Pf*A-M17 inhibitors. The *Pf*A-M17 specific inhibitor exhibited a 22× increase in the IC_50_ value in the resistant line when compared to the parental line, reflecting that its target – a functional *Pf*A-M17 hexamer – was not present, or present at a reduced concentration compared to WT, in keeping with the mechanism of resistance. Whilst the same amount of target is present for compound **3** to bind, it is no longer all in a functional form and as such results in resistance to a *Pf*A-M17 only inhibitor, highlighting the significance of **MMV1557817** as a dual inhibitor. The **MMV1557817** resistant parasites also showed an apparent reliance on *Pf*A-M1, seen by the sensitisation to the *Pf*A-M1 specific inhibitor. Although these aminopeptidases have been shown to have different substrate preferences (15), *Pf*A-M1 is able to cleave a subset of *Pf*A-M17 residues but cannot account for its complete loss (7). Additionally, loss of *Pf*A-M17 has not previously been shown to result in a build-up of apparent *Pf*A-M17 specific peptides but rather an overall increase in haemoglobin-derived peptides (7). This suggests that any decrease in functional *Pf*A-M17 likely results in an overall perturbation in peptide digestion which may account for the *Pf*A-M1 sensitisation seen in **MMV1557817** resistant parasites. This could also be the driving factor for an overall increase in haemoglobin digestion in an attempt by the parasites to increase the free amino acid pool to overcome partial loss of *Pf*A-M17.

**MMV1557817** resistant parasites were also found to be marginally sensitised to artemisinin in the IC_50_ assay performed here. Whilst this small degree of sensitisation is unlikely to be clinically relevant, it suggests that inhibition of end-stage haemoglobin digestion is an attractive combination for artemisinin and its derivatives, with resistance to either compound less likely if used together. Whether this sensitisation is due to the increase in haemoglobin digestion in these parasites or another stressor is yet to be determined. It could, however, be expected that an increase in haemoglobin digestion may also increase the potency of other antimalarial compounds targeting this pathway. For example, chloroquine and its derivatives cause parasite death by blocking the detoxification of haem (67), and as such more haemoglobin digestion may influence this drugs’ efficacy. Interestingly, piperaquine resistant parasites were also sensitised to dual aminopeptidase inhibition by **MMV1557817** (61). These parasites digest significantly less haemoglobin, highlighting the overall relationship of **MMV1557817**’s activity with haemoglobin digestion, and suggesting inhibition of *Pf*A-M1 and *Pf*A-M17 could be useful in a number of different combination therapies. The parasite reduction ratio was also found to be modest, suggesting that without alteration it would only be suitable in a combination therapy. The parasite clearance time of 86 hours can be attributed to the 72-hour lag phase before maximal killing occurs. These killing parameters are not dissimilar to atovaquone, which also has a considerable lag time until maximal killing occurs and a 99.9% parasite clearance time greater than 3 days (35). However, considering that wash out experiments were performed after only 24 or 48 h of treatment, and resulted in less than 10% parasite survival when parasites traversed a trophozoite stage, this lag phase is unlikely to be detrimental, particularly if used in partnership with other fast acting drugs or developed further.

**MMV1557817** was also found to be effective against *P. falciparum* gametocytes at sub-micromolar concentrations. Activity was noticeably reduced as the parasites matured to later stage / mature gametocytes, which is not surprising, consistent with this compound targeting aminopeptidases that function in haemoglobin digestion which become less prominent as the sexual stages mature. It remains to be seen whether **MMV1557817** is active against other stages of the *P. falciparum* lifecycle, but it appears that *Pf*A-M1 and *Pf*A-M17 are transcribed in the mosquito stage of the lifecycle (68). *Pf*A-M1 and *Pf*A-M17 digest proteins that originate from sources in addition to haemoglobin (7, 51), and as such could be active throughout the entire lifecycle, making them an excellent antimalarial target. Of significance, **MMV1557817** showed activity against both *P. falciparum* and *P. vivax ex vivo* parasites within a nanomolar range. There was also a trend towards the compound being more effective against *P. vivax*, however this did not reach significance (p=0.0533). Recombinant assays highlight that **MMV1557817** is most potent against *Pv*-M1, with a *K*_i_ of 19 nM, which is likely contributing to its lower IC_50_ value against *ex vivo P. vivax* compared to *P. falciparum*. An antimalarial compound that has cross-species activity is highly desirable.

The safety profile of **MMV1557817** was explored against human M1 aminopeptidases to assess the likelihood of off-target effects. Human LTA4H, ERAP1 and ERAP2 were not found to be significantly inhibited at biologically relevant concentrations of **MMV1557817**, and neither were several HDAC enzymes, suggesting that despite the hydroxamic acid groups ability to bind metal ions, there are no apparent widespread issues with off-target enzyme activity. Exposure in mice indicated a modest half-life of 4.3 hours, however further PK studies to determine PK parameters and oral bioavailability were not conducted. At an oral dose of 50 mg/kg in mice, unbound concentrations of **MMV1557817** were likely present above the unbound IC_50_ for about 14 h. Despite this limited exposure profile, inhibition of aminopeptidases in the rodent malaria model *P. berghei* was sufficient to substantially reduce parasitaemia, with no significant difference seen when compared to the artesunate positive control. Interestingly, *Pb*-M17 is not essential for parasite survival, however knockout of the gene results in a slow growth phenotype (69). Treatment with a dual M1 and M17 inhibitor of this potency is evidently effective, possibly due to the essentiality of *Pb*-M1 coupled with the very low *K*_i_ inhibition constant for this enzyme.

In conclusion, we have characterized a novel on-target aminopeptidase inhibitor that kills multiple *Plasmodium* species and found it to be a candidate for development as a new lead antimalarial compound. **MMV1557817** was confirmed to be active against multiple stages of *P. falciparum* and encouragingly also kills *P. vivax*, an often-neglected species for antimalarial drug development. Resistance to **MMV1557817** resulted in destabilisation of the *Pf*A-M17 hexamer which is required for enzymatic function and is likely limited by the essentiality of this protein. These results, as well as a lack of cross resistance seen with current resistance mechanisms, strongly suggests that **MMV1557817** and more optimised analogues could be useful as a partner drug for current antimalarials. Inhibition of aminopeptidases involved in the terminal stage of haemoglobin digestion should be considered as an addition to current antimalarial combination therapies.

## ACKNOWLEDGEMENTS

We thank the Medicines for Malaria Venture (MMV) for data generated as part of their network. We also thank Professor David A. Fidock from Colombia University, USA for his generous development and supply of **MMV1557817** resistant parasites characterized in this study. We thank Sibylle Sax from the Swiss Tropical and Public Health Institute, Switzerland for support with the *Plasmodium falciparum* strain panel assays. We acknowledge the Australian Synchrotron for the use of their facility and the Australian Red Cross Lifeblood for providing the red blood cells used in this study.

RCSE, TRM and KD were recipients of an Australian Government Research Training Program Stipend. SD was supported by a NHMRC Dora Lush Biomedical Postgraduate Scholarship (AP1150359), Griffith University DVCR Postgraduate Top-up Scholarship and a Discovery Biology Top-up Scholarship. The Centre for Drug Candidate Optimisation is partially supported by Monash University and Therapeutic Innovation Australia (TIA) through the Australian Government National Collaborative Research Infrastructure Strategy (NCRIS) program. This work was supported by an NHMRC Synergy Grant (1185354) and TFdK-W was supported by an NHMRC Fellowship (1136300).

## Supplementary Data

**Supplementary Figure 1.**
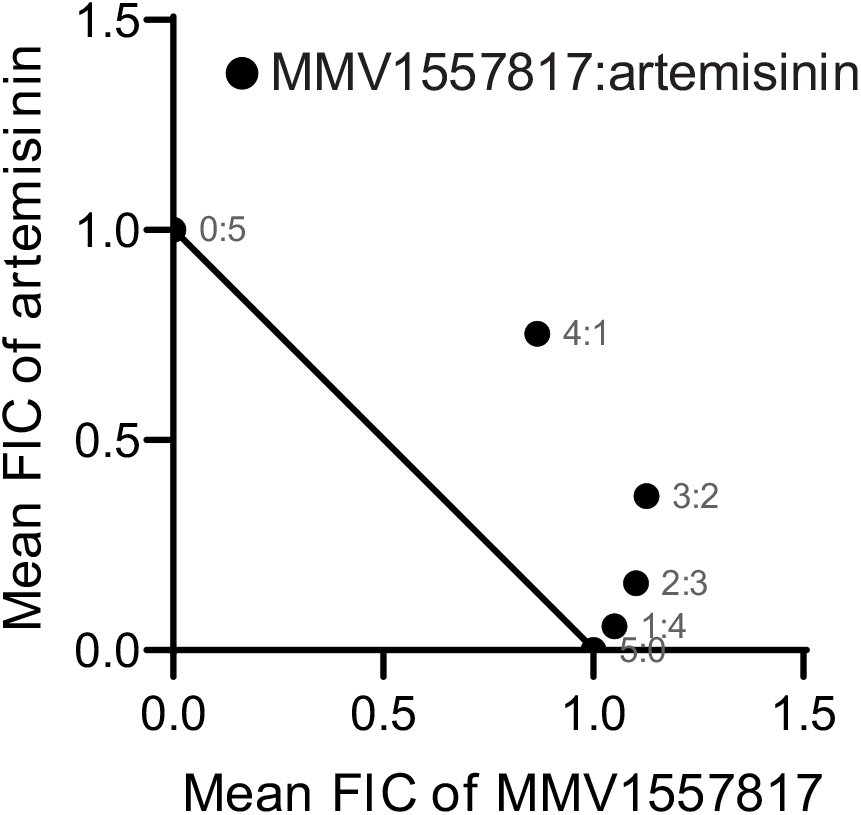
Isobologram showing the interaction between **MMV1557817** and artemisinin tested in Dd2 parasites. Fractional inhibitory concentration (FIC) ≤2≥1 indicate additive effects of the individual compounds. n = 3 performed in triplicate.

**Supplementary Figure 2.**
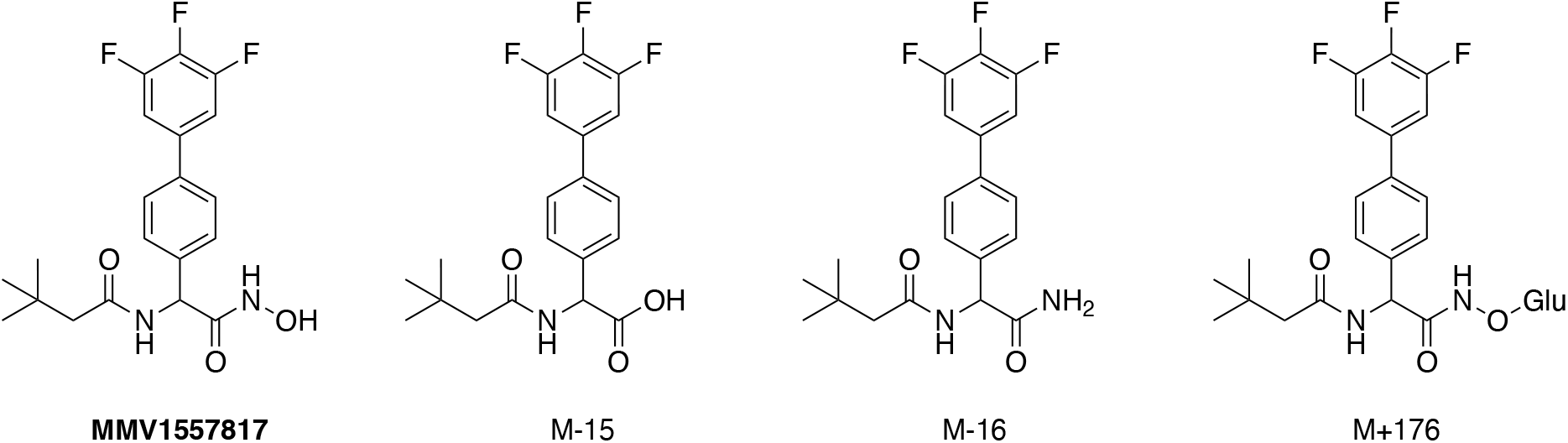
MMV1557817 and putative metabolites detected following incubation with human and rat cryopreserved hepatocytes.

**Supplementary Figure 3.**
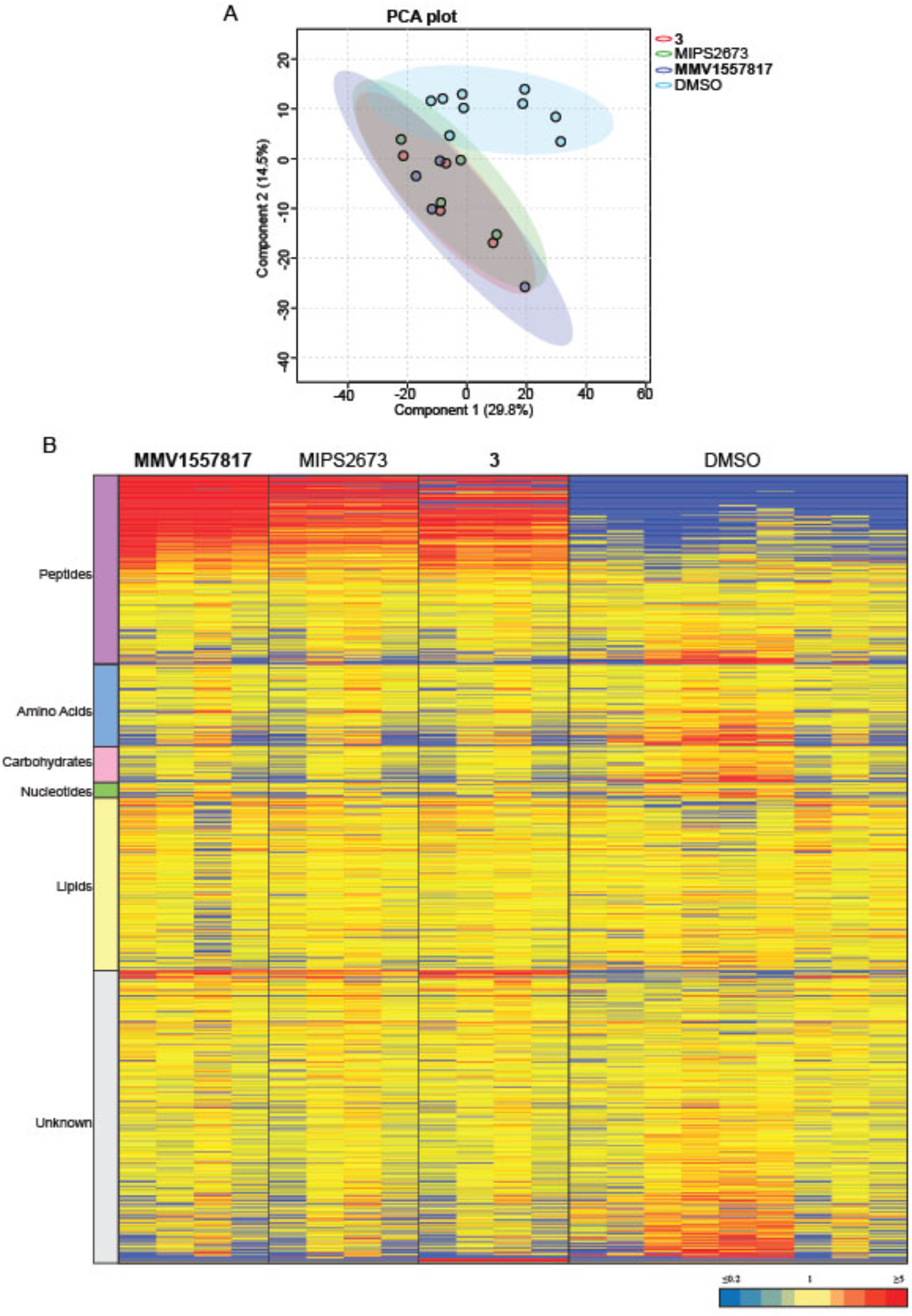
Untargeted metabolomics analysis of *Pf*3D7 parasites treated with **MMV1557817**, MIPS2673 (*Pf*A-M1 inhibitor), **3** (*Pf*A-M17 inhibitor), and DMSO control. (A) Principal component analysis (PCA) of parasites treated with MIPS1557817, MIPS2673 (*Pf*A-M1 inhibitor (51)), **3** (*Pf*A-M17 inhibitor (7)), and DMSO control. Scores plot shows principal components one and two, data points indicate individual sample replicates within each condition and the shaded area denotes 95% confidence interval. (B) Heatmap analysis of peak intensities of all putative metabolites for each condition. Data is shown from 4-7 biological replicates, red, blue and yellow indicates increase, decrease and no change respectively in the relative abundance of putative metabolites identified. Data for **3** and MIPS2673 has been previously published (7, 51).

**Supplementary Figure 4.**
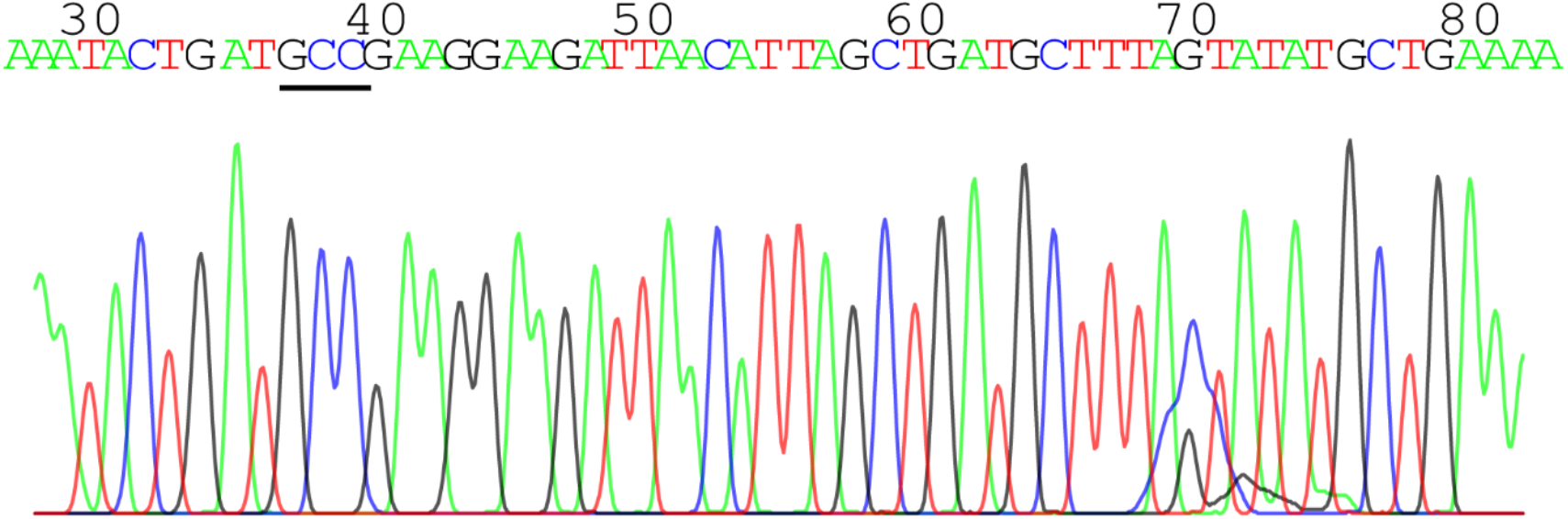
Chromatogram of Sanger sequencing of DNA extracted from parasites harbouring a silent mutation (gct to gcc; underlined in black) at A460 in *Pfa-m17* introduced using CRISPR-Cas9.

**Supplementary Figure 5.**
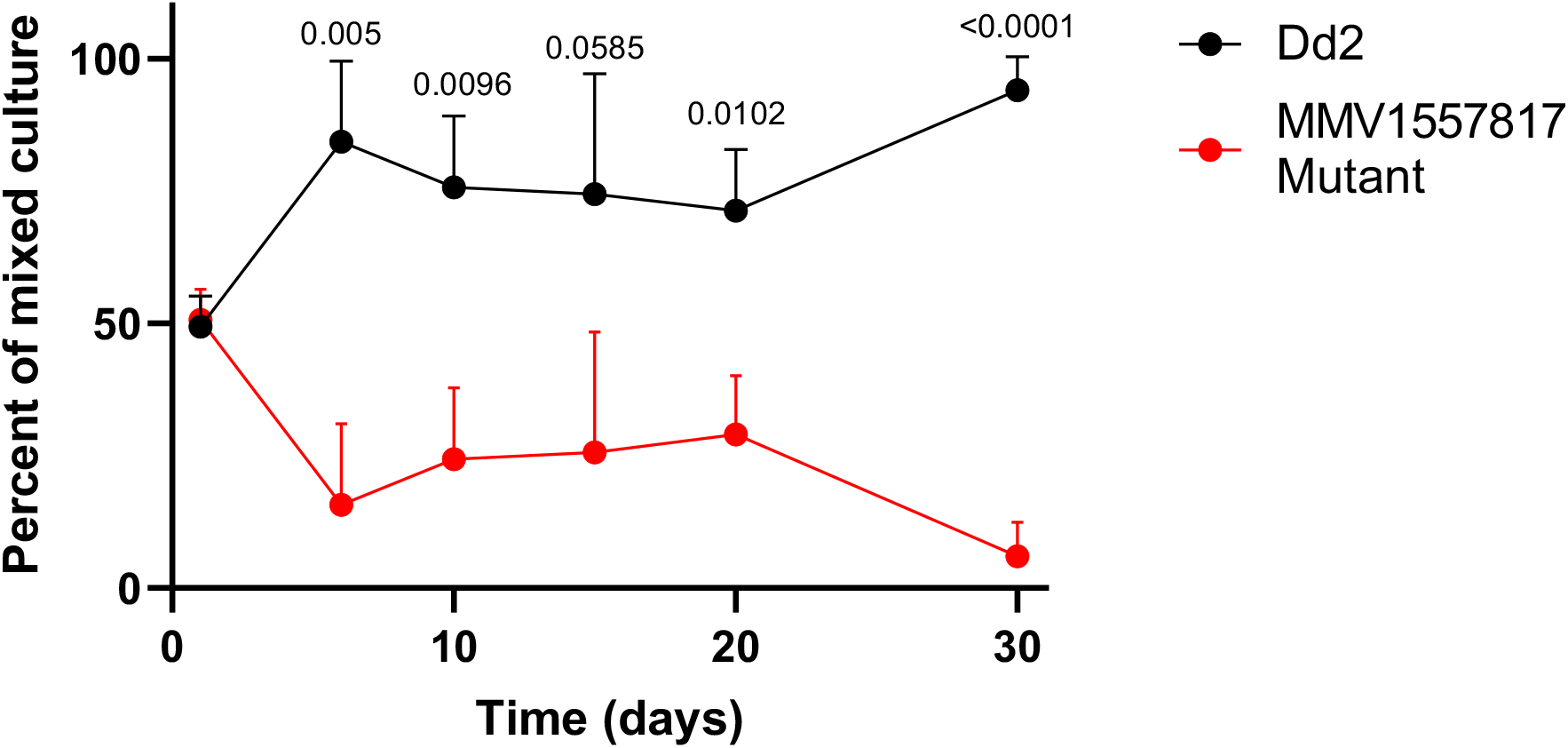
Growth competition analysis of **MMV1557817** resistance parasites versus Dd2 as identified through SNP identification by Sanger Sequencing of the A460 locus of *pfa-m17*. Statistical significance indicated above and determined for each timepoint using an unpaired t-test, n = 3.

**Supplementary Figure 6.**
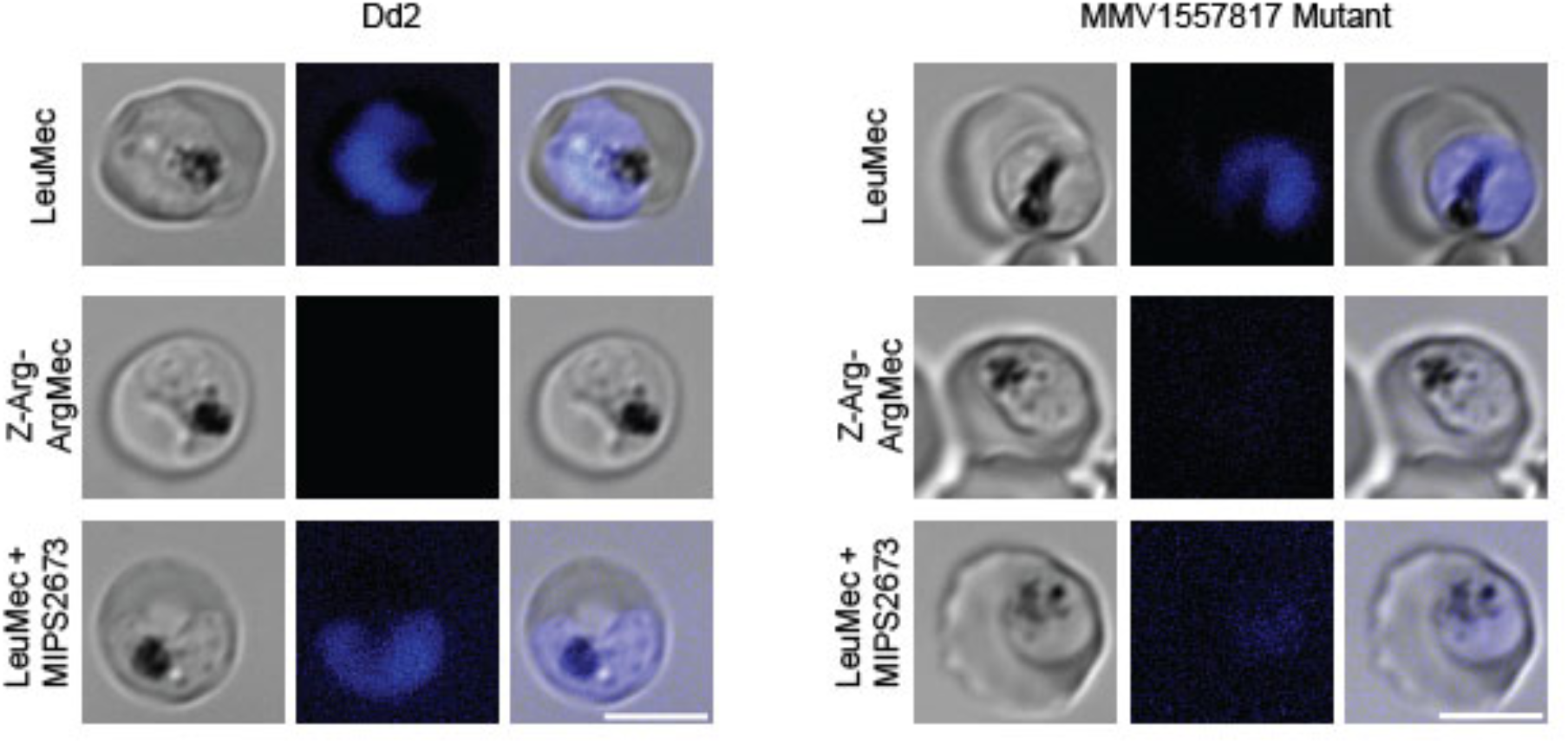
Representative images of live cell Dd2 and MMV1557817 resistant parasites after treatment with the indicated fluorescent substrates. The bottom panel has additionally been treated with the *Pf*A-M1 specific inhibitor MIPS2673 at 10× IC_50_ (3.2 µM; (51)). Scale bar 5 µM.

**Supplementary Table 1:** Data collection and refinement statistics for X-ray crystal structures.

|  | *PfA-M1 - MMV1557817* | *PfA-M17 - MMV1557817* | *PfA-M17(A460S)* |
| --- | --- | --- | --- |
| PDB ID | 8SVL | 8SVM | 8SW9 |
| *Data Collection* |  |  |  |
| Wavelength | 0.95370 | 0.95370 | 0.95370 |
| Resolution range | 41.29 - 1.504 (1.56 - 1.50) | 46.5 - 2.3 (2.382 - 2.3) | 49.57 - 2.6 (2.69 - 2.60) |
| Space group | P 21 21 21 | P 21 21 21 | P 21 21 21 |
| Unit cell (a, b, c,<br>α, β, γ) | 74.90 109.01 118.03<br>90 90 90 | 174.20 177.93 229.76<br>90 90 90 | 174.34 176.73 225.21<br>90 90 90 |
| Total reflections | 1055651 (47886) |  | 425739 (42117) |
| Unique reflections | 153386 (15196) | 306753 (30690) | 212977 (21059) |
| Multiplicity | 6.9 (6.4) | 3.0 (2.9) | 2.0 (2.0) |
| Completeness (%) | 99.97 (99.97) | 96.59 (94.09) | 99.70 (99.91) |
| Mean I/sigma(I) | 11.5 (1.1) | 3.0 (1.5) | 3.88 (1.17) |
| Wilson B-factor | 16.10 | 24.57 | 34.69 |
| R-pim (all I+ & I-) | 0.042 (0.685) | 0.223 (1.483) | 0.1162 (0.6269) |
| CC1/2 | 0.999 (0.431) | 0.931 (0.150) | 0.989 (0.641) |
| *Refinement statistics* |  |  |  |
| Reflections used in refinement | 153367 (15192) | 303998 (29370) | 212390 (21040) |
| Reflections used for R-free | 7575 (705) | 15065 (1432) | 10830 (1062) |
| R-work | 0.1566 (0.2736) | 0.2068 (0.3059) | 0.2348 (0.3156) |
| R-free | 0.1886 (0.2933) | 0.2529 (0.3713) | 0.2804 (0.3584) |
| Number of non-hydrogen atoms | 8676 | 50865 | 48047 |
| macromolecules | 7310 | 47076 | 46600 |
| ligands | 82 | 756 | 383 |
| solvent | 1284 | 3033 | 1064 |
| Protein residues | 889 | 6177 | 6150 |
| RMS (bonds) | 0.019 | 0.004 | 0.002 |
| RMS (angles) | 1.69 | 0.72 | 0.59 |
| Ramachandran favored (%) | 98.20 | 96.77 | 96.52 |
| Ramachandran allowed (%) | 1.69 | 3.13 | 3.08 |
| Ramachandran outliers (%) | 0.11 | 0.10 | 0.39 |
| Rotamer outliers (%) | 0.37 | 0.90 | 2.82 |
| Clashscore | 3.69 | 3.53 | 6.22 |
| Average B-factor | 22.77 | 27.02 | 39.66 |
| macromolecules | 20.27 | 26.39 | 39.65 |
| ligands | 31.19 | 43.59 | 50.38 |
| solvent | 36.44 | 32.54 | 36.02 |

Statistics for the highest-resolution shell are shown in parentheses.

**Supplementary Table 2:** Interaction between artemisinin and MMV1557817 against *P. falciparum* Dd2 parasites.

| Ratio (MMV1557817:<br>artemisinin) | MMV1557817<br>mean FIC ± SEM | artemisinin mean<br>FIC ± SEM | $\Sigma$ FICs<br>(interaction) |
| --- | --- | --- | --- |
| **0:5** | 0 | 1.00 ± 0.12 | |
| **1:4** | 0.87 ± 0.08 | 0.75 ± 0.07 | 1.62 (Additive) |
| **2:3** | 1.13 ± 0.19 | 0.37 ± 0.06 | 1.49 (Additive) |
| **3:2** | 1.10 ± 0.20 | 0.16 ± 0.03 | 1.26 (Additive) |
| **4:1** | 1.05 ± 0.18 | 0.06 ± 0.01 | 1.11 (Additive) |
| **5:0** | 1.00 ± 0.19 | 0 | |

FIC: fractional inhibitory concentration

**Supplementary Table 3:** Baseline characteristics of isolates for which *ex vivo* assay was accomplished.

| Baseline characteristics | *P. falciparum*<br>n=15 | *P. vivax*<br>n=8 |
| --- | --- | --- |
| Total number of isolates reaching harvest (%) | 14 (93) | 6 (75) |
| Median (range) duration of assay (hours) | 44 (41-46) | 47 (44-48) |
| Geometric mean (95% CI<sup>a</sup>), parasitaemia (asexual parasites/μL) | 31,779 (14,914-67,713) | 9,837 (3,124 – 30,975) |
| Median initial % (range) of parasites at ring stage | 100<sup>b</sup> | 90 (76-99) |
| Mean (95% CI) schizont count at harvest | 50(34-86) | 45 (28-55) |

<sup>a</sup> CI, confidence interval
<sup>b</sup> No range given (all values were 100%)

**Supplementary Table 4.**
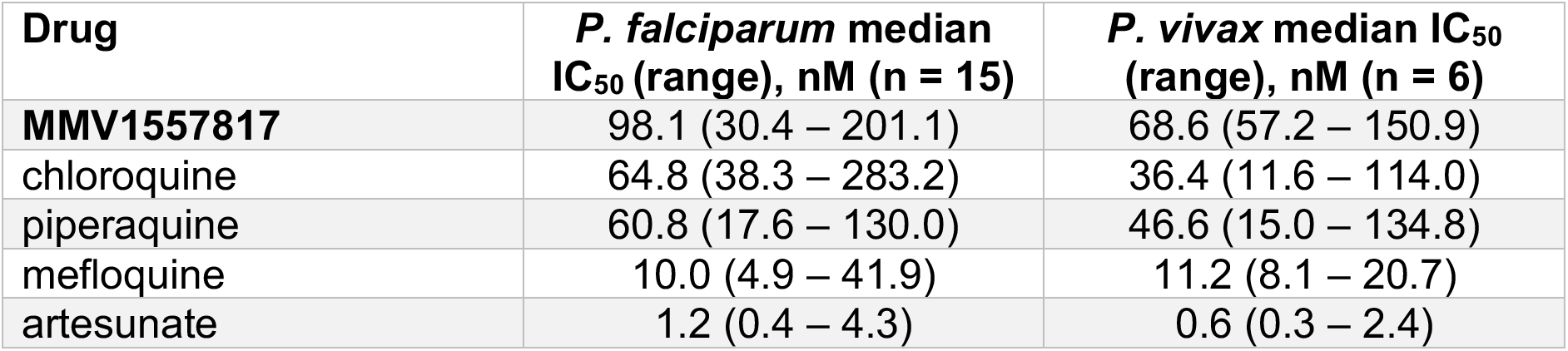
*Ex vivo* drug susceptibility to MMV1557818 and standard anti-malarials according to species tested.

| **Drug** | ***P. falciparum* median IC<sub>50</sub> (range), nM (n = 15)** | ***P. vivax* median IC<sub>50</sub> (range), nM (n = 6)** |
| --- | --- | --- |
| **MMV1557817** | 98.1 (30.4 – 201.1) | 68.6 (57.2 – 150.9) |
| chloroquine | 64.8 (38.3 – 283.2) | 36.4 (11.6 – 114.0) |
| piperaquine | 60.8 (17.6 – 130.0) | 46.6 (15.0 – 134.8) |
| mefloquine | 10.0 (4.9 – 41.9) | 11.2 (8.1 – 20.7) |
| artesunate | 1.2 (0.4 – 4.3) | 0.6 (0.3 – 2.4) |

**Supplementary Table 5:** Percent inhibition of control enzyme activity.

| **Protein** | **Mean % inhibition by MMV1557817 (n=2)** |
| --- | --- |
| HDAC1 | 2.8 |
| HDAC2 | 3.0 |
| HDAC5 | -2.8 |
| HDAC7 | 7.7 |
| HDAC8 | 6.3 |
| HDAC9 | -3.6 |
| HDAC10 | 5.3 |
| MMP-2 | 39.8 |
| MMP-3 | -0.8 |
| MMP-7 | -6.7 |
| **MMP-8** | **52.6** |
| MMP-9 | 46.3 |
| MMP-14 | 24.1 |

